# Measuring the threat from a distance: insight into the complexity and perspectives for implementing sentinel plantation to test host range of *Xylella fastidiosa*

**DOI:** 10.1101/2022.07.22.500186

**Authors:** Noemi Casarin, Séverine Hasbroucq, Júlia López-Mercadal, Miguel Ángel Miranda, Claude Bragard, Jean-Claude Grégoire

**Affiliations:** Earth and Life Institute Applied Microbiology (ELIM), Université catholique de Louvain (UCLouvain), Croix du Sud 2 bte L7.05.03, 1348 Louvain-la-Neuve, Belgium; Spatial Epidemiology lab (SpELL), Université libre de Bruxelles (ULB), CP 160/12, 50 av. F.D. Roosevelt, 1050 Bruxelles, Belgium; Zoologia Aplicada i de la Conservació (ZAP), Universitat de les Illes Balears (UIB), Cra. De Valldemossa, km 7.5, 07122 Palma de Mallorca, Illes Balears, Spain

**Author notes:** Correspondence: Jean-Claude Grégoire,; Claude Bragard.

**Keywords:** biological invasions, *ex-patria* planting, Majorca, northern Europe, pest risk analysis, *Prunus domestica*, *Quercus petraea*, *Salix alba*

## Abstract

The sentinel plantation concept consists of assessing the impact of exotic factors, such as pests and pathogens, on plants of interest by planting them out of their native range. This tool is a way to enhance knowledge for pest risk analysis (PRA) by guiding decisions on how quarantine organisms should be regulated and where to focus prevention and surveillance efforts for an early detection. In this study, the sentinel method was used in the case of research on *Xylella fastidiosa*, a plant pathogenic bacterium that has recently been found established in southern Europe, but whose potential impact and possible host range are still poorly documented in northern areas where the bacterium is not known to occur. To improve knowledge on the susceptibility of potential hosts of *X. fastidiosa* in northern Europe, a sentinel plantation of *Prunus domestica* cv. Opal, *Quercus petraea* and *Salix alba* was established in the *X. fastidiosa*-infected area of Majorca. In order to assess the circulation of the bacterium in the sentinel plot and around it, surveys of the local flora and insect vectors were carried out, as well as the planting of a network of rosemary “spy plants”. Symptomatic monitoring and molecular analyses were performed on the sentinel plants for four years. During these years, *X. fastidiosa* was never detected in our sentinel plants most likely because of the low infectivity pressure recorded in the surroundings. This study underlines the complexity of conducting sentinel plantation assays combined with *X. fastidiosa* research, highlighting the need for long-term investigation and questioning the efficiency of the sentinel tool. However, this study is placed in perspective with other valuable sentinel plantations. It also highlights the complementarity of the tool and proposes elements to improve or reorient the implementation of future sentinel projects.

## Introduction

The world sustainability is threatened by outbreaks of invasive pests and pathogens increasingly spreading around the globe (Simberloff et al. 2013; Diagne et al. 2021). These organisms largely travel to new areas through global trade, with living plants or with wood packaging material, which are considered as the main pathways of plant-related organism introductions (Kenis et al. 2007; Liebhold et al. 2012; Santini et al. 2013; Meurisse et al. 2019). These agents often expand by outcompeting native species because they are transported far from their natural enemies (“enemy release hypothesis”; Keane and Crawley 2002; Colautti et al. 2004), allowing them to allocate resources to growth and fecundity instead of defense, enhancing their fitness (“evolution of increased competitive ability” hypothesis; Blossey and Notzold 1995; Manfredini et al. 2013). They may trigger epidemics, sometimes on new hosts whilst they were less harmful to their native hosts, as they have not co-evolved with the new local plants that lack of specific defense mechanisms (Pimentel et al. 2001; Aukema et al. 2011). Apart from trade and globalization, climate change and intensive land-use are also factors enhancing outbreaks by decreasing the resilience of the agricultural production systems and of forests (Walther et al. 2009; Bosso et al. 2016).

Preventing the introduction and the establishment of pests and pathogens in new areas is the most efficient tool for mitigating the consequences of a disease in terms of cost, biodiversity conservation and human impact (Barham et al. 2016). This includes the implementation of a pest risk analysis (PRA), which is an assessment giving biological, scientific and economic information on a particular organism (Aukema et al. 2011; Tomoshevich et al. 2013; EFSA PLH Panel 2018) to understand its potential impact and how it should be regulated (Parker et al. 1999; EU 2000; Liebhold et al. 2012). If considered harmful, the first measure taken to avoid its introduction might be its inclusion in a quarantine list implying either thorough inspections of imported plants before or after the importation, plant production in pest-free areas or sites of production, or complete prohibition of trade or production of its native host plants (EU 2000).

However, these measures are not fully effective by themselves. Inspections can fail to intercept all the potential pests and pathogens travelling through plant trade (Kenis et al. 2007; Eschen et al. 2015, 2017, 2019). First, these agents can be invisible to the naked eye because of their intrinsic nature or because there are in a latent form or in an endophytic stage on their traded hosts, leading to asymptomatic infections (Stergiopoulos and Gordon 2014; Migliorini et al. 2015). Secondly, despite the prioritization of inspected organisms through PRA, the massive volume of traded materials makes the systematic control of each plant inoperable, only batches will be thoroughly examined (Britton et al. 2010; Eschen et al. 2015). Finally, PRA relies on prior awareness and knowledge of a pest and this knowledge is not always available; several agents, including non-catalogued taxa, harmless in their native region, are unknown to be invasive and pathogenic prior their introduction in a new land, and escape controls (Brasier 2008; Britton et al. 2010; Tomoshevich et al. 2013). The few of them that manage to establish and cause significant damage are then often discovered too late to avoid outbreaks. Such is the case for some of the most damaging organisms of temperate forests that have occurred in recent years, which were unknown as pests prior to their introduction in a new area (Britton et al. 2010). Examples are the epidemics of Dutch elm disease caused by *Ophiostoma ulmi* Buisman and *O. novo-ulmi* Brasier that decimated billions of elm trees in Europe and America in the 20^th^ century (Brasier and Buck 2001), or the massive damage to pines in Asia (Zhao et al. 2008) and Europe (Soliman et al. 2012) caused by the pine wood nematode, *Bursaphelenchus xylophilus* (Steiner and Buhrer 1934) Nickle, 1970, an organism well tolerated by its native pine hosts in North America (Akbulut and Stamps 2012).

A way to enhance knowledge about potentially damaging organisms to improve biosecurity systems would be to expose plants of interest out of their native range to study their susceptibility to local organisms in specific relevant locations, e.g. a frequent plant exporting country (Roques et al. 2015; Barham et al. 2016). These plants would represent sentinels for their species in the foreign land. They provide an early warning for potential threats and additional information for PRA to set preventing measures and to know where the efforts for plant protection should be focused (Barham et al. 2016; Mansfield et al. 2019). An EPPO standard document was published in 2020, “PM 3/91 Standard on Sentinel woody plants” (EPPO 2020), to explain the approach and to provide guidance to carry out sentinel plant studies to identify new pest risks.

Sentinel plant research can be carried out by different ways (Britton et al. 2010). A first way is through botanical gardens and arboreta gathering a collection of specimens from all over the world, which are generally out of their area of origin and exposed to local agents. For such studies, the International Plant Sentinel Network (IPSN), working closely with National Plant Protection Organizations (NPPOs), was created. It connects the botanical gardens and arboreta staff around the world and gives them tools and expertise to monitor and to identify new pests and pathogens (Barham et al. 2016). Tomoshevich et al. (2013) for example, discovered 29 new pest-host associations whose 18 noticeably damaging for European trees by studying European and Eurasian trees in Siberian gardens in Russia. However, in botanical gardens and arboreta, the number of representatives of each plant species is generally limited (Roques et al. 2015), the trees are often large and hence difficult to examine in detail, and they are usually subject to pesticide treatments or other management practices, which ensure plant health in the gardens (Eschen et al. 2019). Furthermore, gardens are often located in urban areas distant from the habitats of potential pests. All these reasons reduce the likelihood for an organism to reach and infect a specific plant species in an arboretum (Britton et al. 2010). A second way to conduct sentinel plant researches is directly establishing actual plantations of exotic plants of interest in an environment where we want to study the impact of local pests and pathogens, the so-called “sentinel plantations” (Roques et al. 2015) or *“ex-patria* plantings” (Eschen et al. 2019). For example, Roques et al. (2015) and Vettraino et al. (2015) established two sentinel plantations of European tree species in China to investigate new pest-host associations potentially threatening to Europe that may emerge as a result of trade.

On the other hand, some well-known pathogens are still restricted to one part of the world and their potential host range in non-infected areas is uncertain and must be investigated. Such is the case of the phytopathogenic bacterium *Xylella fastidiosa* Wells et al., 1987, with more than 650 reported host plant species, and for which the host range continues to extend as the bacterium enters new areas (EFSA 2022). While the threat of *X. fastidiosa* is definite for the European Mediterranean flora, the potential impact for northern areas is uncertain as most of the flora in these regions has never been exposed to the bacterium and probably contains many unreported hosts. The objective of our study was therefore to establish a sentinel plantation with European northern trees in a *X. fastidiosa* infected area in order to study the potential host range for these still-uninfected regions.

The gammaproteobacterium *X. fastidiosa* (Xanthomonadaceae) is strictly limited to the foregut of xylem sap-feeding insect vectors, mainly leafhoppers and spittlebugs (Hemiptera, Cicadomorpha) (Redak et al. 2004; Almeida et al. 2005; Chatterjee et al. 2008) and to the xylem vessels of its host plants. While many listed hosts are asymptomatic, the bacterium causes severe outbreaks on several crops, ornamental plants and shade trees generally provoking leaf-scorching that could lead to plant death (EFSA PLH Panel 2018). First limited to the Americas, the bacterium is currently regulated in Europe as a quarantine organism under the Council Directive 2000/29/EC (EU 2000). Between 2014 and today, the Europhyt database recorded 51 interceptions of *X. fastidiosa* in plants for planting and four interceptions of leafhoppers (EUROPHYT Online database; EFSA PLH Panel 2018). Despite the border controls and EU prevention measures, a first focus of *X. fastidiosa* in Europe was discovered in 2013 in Apulian olive groves (Italy), for which more than 21 million olive trees were estimated to be affected in 2018 (Saponari et al. 2019). The bacterium was then identified in mainland France, in Corsica (Denancé et al. 2017), in mainland Spain, in the Balearic Islands (Olmo et al. 2017), in another region of Italy (Tuscany) (Saponari et al. 2019) and in Portugal (EFSA PLH Panel 2019b; EUROPHYT Online database). Divided into several subspecies (mainly subsp. *fastidiosa*, subsp. *multiplex*, and subsp. *pauca*; Schaad et al. 2004) and more finely according to its sequence type (ST) (Scally et al. 2005; Yuan et al. 2010), 11 different STs were identified throughout Europe revealing multiple independent *X. fastidiosa* introduction events (EFSA 2021; Cunty et al. 2022). Phylogeny studies allowed to date back to the different entries of *X. fastidiosa* in the specific European regions, indicating entrance in the 1980s, 1990s and 2000s according to the area, i.e. well before the official identification of the pathogen’s establishment on the continent. *Xylella fastidiosa* is therefore a perfect example of an organism escaping control due to the complexity of detection given the asymptomatic pool of hosts, the potentially long latent period limiting visual inspection, and the number of reported and supposed-unreported hosts, as well as the lack of specific surveillance programs and the limited availability of specific diagnostic tools in the past. Its movement into Europe has been caused in part by the trade of asymptomatic coffee plants imported from Latin America (EFSA PLH Panel 2015; Denancé et al. 2017).

However, it has been shown that eradication of *X. fastidiosa* may be complex if not impractical once it is well established and has reached a large geographical extent (Strona et al. 2017; EFSA PLH Panel 2019a). Therefore, while entries can hardly be prevented, early detection is primordial to limit damage of outbreaks. The probability of early detection would increase by improving knowledge on where the bacterium is most likely to establish in order to conduct effective surveillance. In fact, performing detection tests on every plant in random areas is neither efficient nor conceivable as it would exceed any diagnostic capability considering the wide range of potential host plants (EFSA PLH Panel 2016). Targeting the main host plants and establishing a prioritization list, is essential to know where to focus resources and monitoring efforts.

To enhance knowledge on the susceptibility of potential hosts of *X. fastidiosa* in northern Europe, a sentinel plantation of northern plant species *Prunus domestica* cv. *Opal*, *Quercus petraea* and *Salix alba*, was established in the *X. fastidiosa-infected* area of Majorca (Balearic Islands, Spain). There, the bacterium is considered widespread and well established. Three different STs belonging to two subspecies (*X. fastidiosa* subsp. *fastidiosa* ST1, and *X. fastidiosa* subsp. *multiplex* ST81 and ST7) have been identified on several hosts including wild olives, cultivated olives, almonds, grapes and figs (Olmo et al. 2021). They are mainly transmitted by the *Philaenus spumarius* Linnaeus, 1758 (Aphrophoridae) vector and to a lesser extent, by *Neophilaenus campestris* Fallén, 1805 (Aphrophoridae) (López-Mercadal et al. 2021). This study experiments the sentinel plantation tool in the case of *X. fastidiosa* research. The outcome questions the efficiency of the method, at least in this particular case, and highlights the complexity of its implementation. However, it provides a methodology and several perspectives for future sentinel projects.

## Methods

### Preliminary tests and plant movement

The establishment and the monitoring of the sentinel plantation was achieved with the collaboration of the ZAP group of the University of the Balearic Islands (UIB). First, the agreement of the local government and the UIB authorities had to be obtained. Then, the plant material was bought at the Calle-Plant nursery in Wetteren, Belgium. It consisted in dormant material: 30 *Salix alba* 0/1 80/120, 30 *Quercus petraea* 2/0 80/100 and 30 *Prunus domestica* cv. Opal 2 years grafted on Myrobolan or St Julien. Although all the plants were equipped with a phytosanitary certificate, *X. fastidiosa* specific detection tests were performed on several twigs of each plants to make sure the initial material was free of the bacterium. For this purpose, three branch parts of each plant were collected and bark-peeled. They were chopped and their DNA was extracted according to the CTAB-based DNA extraction protocol specific for *X. fastidiosa* plant samples (“PM 7/24 (4) *Xylella fastidiosa*”, EPPO 2019). The detection was then performed by PCR (Minsavage et al. 1994). After this double check, the ninety plants were wrapped in hessian bags filled with wood chips and were brought by truck from Belgium until the UIB campus in Palma (Majorca, Balearic Islands) on March 2018. The chips were humidified during the 2-day trip to avoid root dryness.

### Location and establishment

The location of the plot was chosen with the UIB collaborators mainly based on the ease of connection to an irrigation system, as well as on the observation of *Philaenus spumarius* and *Neophilaenus campestris* nymphs on the ground vegetation and the presence of host plants such as wild olive and almond trees. For the positioning of the plants in the plot, the JMP^®^ software was used to generate nine blocks, each one composed by three plants of each species randomly distributed (Fig. 1). The scheme was divided by blocks to take into account the potential gradients such as the slope, irrigation distribution, or sunlight. The trees were planted directly into the ground to promote the growth of the root system and to enable them to survive throughout the season (Fig. 2). The soil was compact and rocky and was dug thanks to machines (Fig. 3). In every hole, about 20 liters of breeding soil were poured. The trees were separated from each other by 1.50 m and the whole plantation covered a total area of 12 m^2^. The irrigation system was established in the second year of the plantation. It consisted in three closed loops of pipe with one dripper per plant, allowing a constant pressure in all pipes and the same amount of water per plant (Fig. 1). The climatic data were followed through the season thanks to a HOBO^®^ device placed in the middle of the plantation.

**Figure 1.**
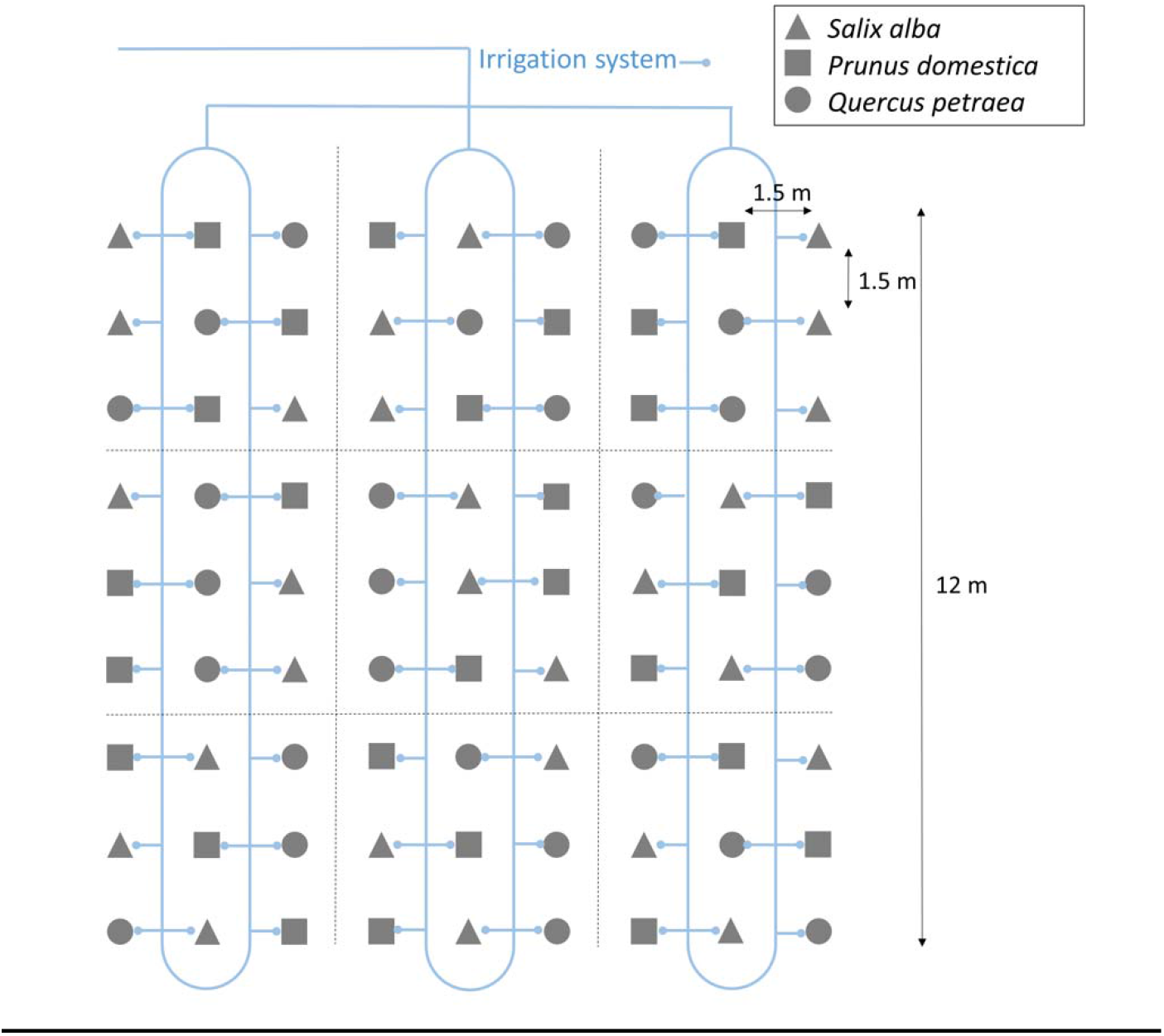
Scheme of the sentinel plantation of *Salix alba, Prunus domestica* cv. Opal and *Quercus petraea*. The dotted lines delimit nine blocks in which there are three plants of each species distributed randomly (JMP^®^). The solid blue line is for the representation of the irrigation system consisting in three closed loops of pipe with one dripper per plant.

**Figure 2.**
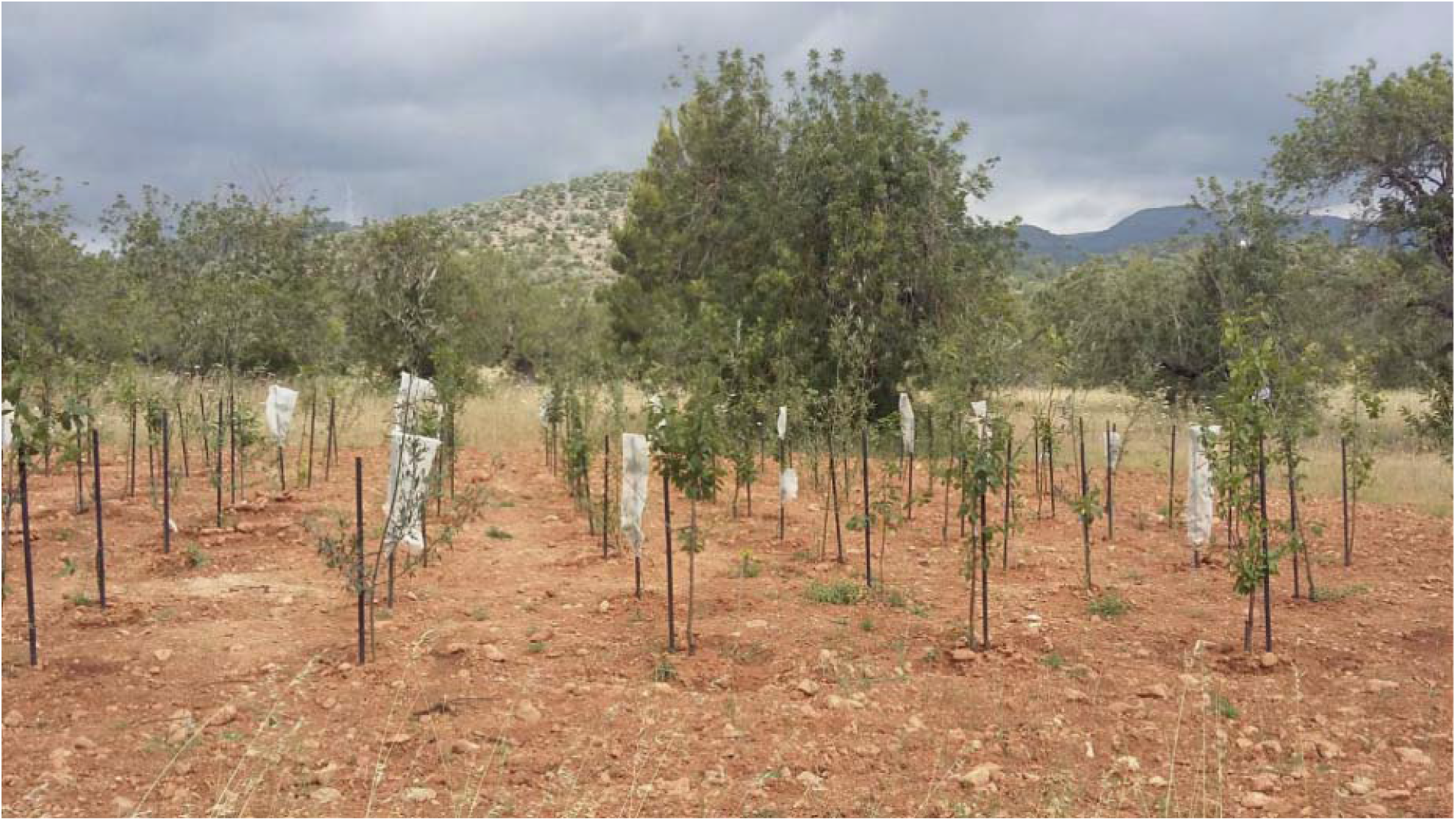
Picture of the Belgian sentinel plantation in Palma (Majorca, Balearic 237 Island) in May 2018.

**Figure 3.**
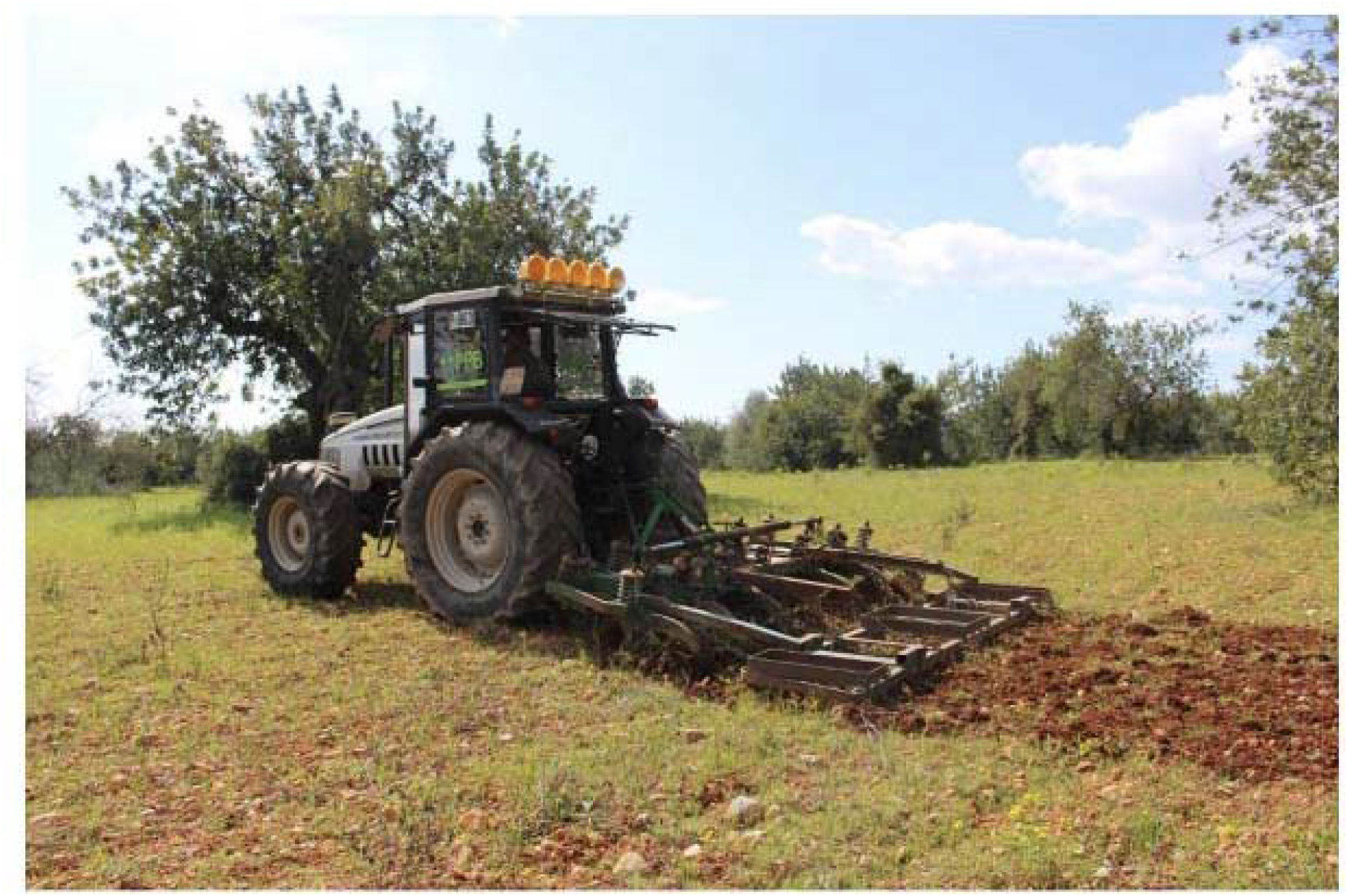
Preparing the ground for planting the sentinel trees. The pictures highlight the difficulty of establishing the sentinel plantation in the dry, compact and rocky soil of the area.

### Exploring the surroundings

To monitor the circulation of the bacterium in the plot and around it, a 100-m demarcated area was organized around the plantation. In this area, i. a floristic inventory was carried out; ii. insect vectors were sampled; iii. a rosemary “spy plant” network was established (Fig. 4).

**Figure 4.**
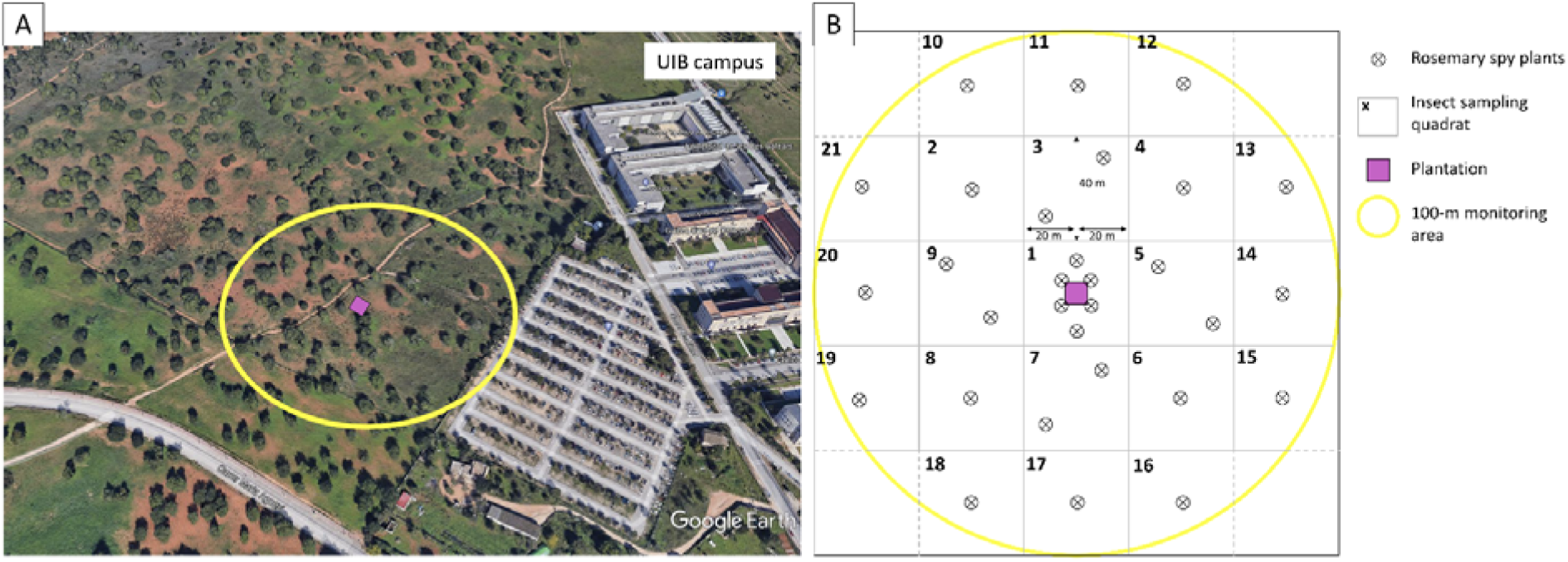
Surroundings of the sentinel plantation. **A.** Google Earth view (Google Earth Pro, satellite image of May 6, 2021) of the UIB campus with the location of the sentinel plantation (purple square) and the 100-m demarcated area around the plantation (yellow circle) **B.** Scheme of the plantation and the demarcated area. In the demarcated area, a floristic inventory was carried out, insect vectors were sampled in determined quadrat and a rosemary “spy plant” network was established by planting evenly seedlings around the plantation.

#### Floristic inventory

To locate and assess the proportion of *X. fastidiosa* host plants in the area and to follow the eventual appearance of symptoms, a floristic inventory was carried out. It consisted in identifying and mapping the tree layer of the demarcated area. An identification of the main herbaceous species was also performed with the help of local collaborators and of a flora determination key handbook.

#### Rosemary network

44 *Rosmarinus officinalis* were planted around the campus: 32 plants evenly positioned in the demarcated area (Fig. 4) and 12 plants in other places of the campus. The idea was to choose a robust plant adapted to local environmental conditions and which is quite susceptible to several subspecies of *X. fastidiosa*. Planting and regularly sampling these susceptible plants for bacterial detection provide a spy network allowing to control the circulation of the bacteria in the vicinity of the plantation. The plants were bought in a local nursery in March 2018. They were first checked for *X. fastidiosa* presence with molecular tests before planting them, consisting of a CTAB-based DNA extraction followed by PCR of Minsavage et al. (1994). For sampling, about 15 leaves were collected on each plant, starting with symptomatic ones, and were processed right away in the local laboratory. The midrib and the petiole were sectioned and the total DNA was extracted with the CTAB-based extraction procedure (EPPO 2019). The DNA samples were then sent to Belgium and were processed at UCLouvain by PCR of Minsavage et al. (1994) in the first three years, and by real-time PCR of Harper et al. (2010) in the fourth year-final testing. In this final year, about five twigs per plant were collected as well and were processed in the same way.

#### Insect sampling and testing

Insects were sampled with two objectives. On the first hand, they were collected to be tested for *X. fastidiosa* presence by PCR (Minsavage et al. 1994) and quantitative PCR (Harper et al. 2010) to check for the circulation of the bacterium around the plantation. On the other hand, during the first year, the vector population density was assessed every month to determine the variability of the potential transmission during the season. For this study, the 100-m area around the plot was divided in 25 blocks (Fig. 4). In each block, the same number of insect samples were undertaken. According to the development stage of the insect, the sampling method was adapted. For the nymphal stage, a frame of 50 cm x 50 cm was used (0.25 m^2^) and was thrown randomly four times in each block. The nymphs present in the surface delimited by the frame were counted. In total, 84 samples were undertaken throughout the demarcated area and the number of nymphs/m^2^ could be estimated. Although *X. fastidiosa* is lost after every molt, the nymphs can also get infected with it (Purcell and Finlay 1979; Redak et al. 2004). Therefore, in addition to the density study, three nymphs of *P. spumarius* and three nymphs of *N. campestris* were collected in each block for bacterial detection to potentially already have an indication of the circulation of the bacteria in the plot. This number was chosen in order to not affect the vector abundance around the plot for the rest of the season.

Regarding insects at adult stage, the sampling was carried out with sweeping nets. Two samples per block were undertaken in the ground layer, one sample corresponding to ten sweepings. The sweepings were done homogeneously in each block in order to cover all the area. In total, 42 samples were undertaken throughout the demarcated area and the number of adult/swept was measured. Again, only three insects per species (*P. spumarius* and *N. campestris*) were collected per block. Due to the small number of insects found in summer, the tree layer was also sampled. All the wild olive, almond and carob trees in the demarcated area were hit fifteen times with sweeping nets, distributed evenly on the plant in order to cover its entire attainable foliage surface. The number of adult/tree could be assessed.

The insects collected were placed at −20 °C, then stored in ethanol 70 % and were sent to Belgium where they were processed. The eyes were removed and the DNA of the head together with the mouthparts was extracted using the CTAB-based protocol (EPPO 2019). The extracted DNA was then processed by PCR of Minsavage et al. (1994), by nested PCR of Cruaud et al. (2018) or by quantitative PCR of Harper et al. (2010).

### Sentinel plantation monitoring

Visual inspections were carried out for each sentinel tree. The appearance of *Xylella*-like symptoms was cautiously observed and wilting, shoot dieback, desiccation, defoliation or any change in leave color were reported. The evolution of the size of the different plants was also monitored, as well as the presence of *Xylella*-vectors or of other pests or organisms. In parallel, molecular analyses were performed on each plant. One sample per plant was collected, consisting of ten leaves per plant and 4-5 small twigs collected from all sides of the plant, but prioritizing symptomatic areas if there were any. DNA extractions were carried out with the CTAB-based protocol (EPPO 2019) on leaf midribs, on petioles and on the twigs after bark peeling and cutting them into small pieces. The DNA samples were then sent to Belgium where they were processed by PCR of Minsavage et al. (1994) in the first three years. In the final-testing of the fourth year, two samples per plants were collected, one sample consisting of 10 different twigs distributed throughout the plant together with 10 to 20 leaves, always prioritizing symptomatic parts. After extraction, they were processed by PCR of Minsavage et al. (1994) as well as by real-time PCR of Harper et al. (2010). No fertilizer was applied and no pruning was carried out in the winter, to allow the plants to develop naturally and not to cut potentially infected sections.

### Sowing ground vegetation

Because the planting of the sentinel plants with machines had removed the herbaceous layer in the sentinel plantation, which could prevent insects from reaching the trees, it was decided resowing grass in February of the second year to reconstitute this layer. The seed consisted of a universal mix of Asteraceae, Fabaceae and Poaceae.

### Monitoring of the plantation for four years

The planning of the plantation monitoring during the four years is available in Table 1. The first year, it was decided to monitor the plantation and the demarcated area almost every month of the vector-season to assess the vector density fluctuation and to measure the rate of infection, if any, of the different plant species. In March, nymphs were sampled while from May to October, insect adults were monitored. Several rosemary plants had desiccated already in May of the first year. Therefore, the dead ones were replaced in May and also in February of the following year. From the second year onwards, the sampling periods were chosen to correspond more or less to the beginning and the end of the highly infectious period of *X. fastidiosa* carried by the insect vectors, respectively June and October, avoiding the estivation periods of insects. The third year was impacted by the Covid-19 crisis and only one sampling campaign could be carried out in October 2020.

**Table 1.**
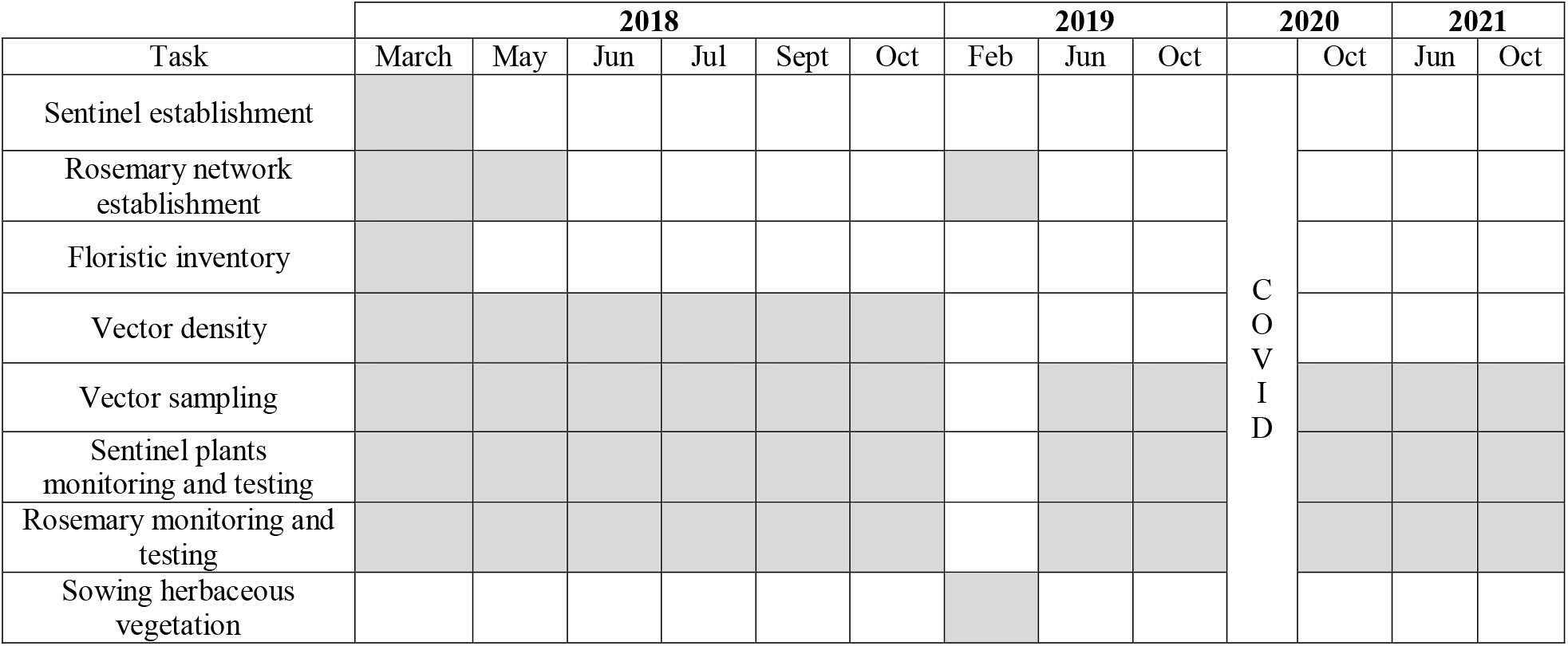
Four-year schedule of the establishment and monitoring of the plantation and of the demarcated area.

## Results

### Insight into surrounding plants

About 170 trees were inventoried: 134 carob trees, 31 wild olive trees, 5 almond trees and 2 pine trees. Their distribution can be observed at the Fig. 5. The wild olive trees and the almond trees are both host plants of *X. fastidiosa*. Therefore, 36 host plants of the bacterium were identified in the 100 m around the plot. Among these host plants, 64% showed typical leaf scorching symptoms of *X. fastidiosa*. Concerning the ground vegetation, the identified plants were mainly: *Conium maculatum* (Apiaceae), *Foeniculum vulgare* (Apiaceae*), Cichorium intybus* (Asteraceae), *Dittrichia viscosa* (Asteraceae), *Galactites tomentosa* (Asteraceae), *Euphorbia medicaginea* (Euphorbiaceae) and many Poaceae (*Oryzopsis* sp. and others).

**Figure 5.**
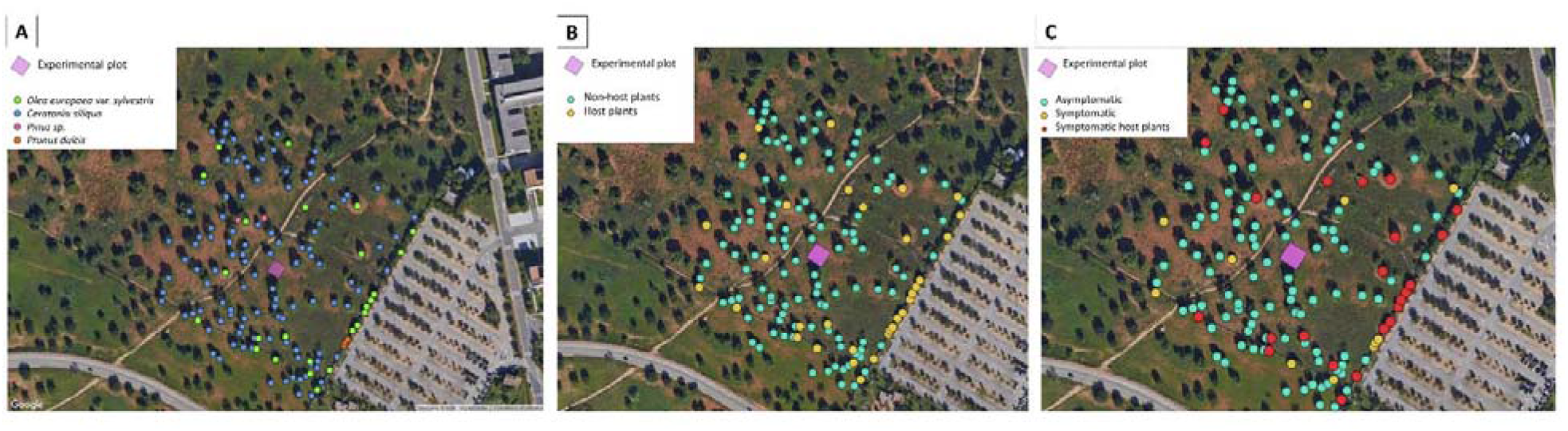
Tree species inventory, host and health status around the sentinel plantation. **A.** Map of the different tree species in the area of 100-m around the plantation. The pink square is the experimental plot (the sentinel plantation). The green dot are for *Olea europaea* var. *sylvestris* (wild olive tree), the blue dot for *Ceratonia siliqua* (carob tree), the pink dot for the *Pinus* sp. (pine tree) and the orange dot for the *Prunus dulcis* (almond tree). **B.** Map of the host status of the trees located in the area of 100-m around the plantation. The green dots are the non-host plants of *X. fastidiosa* and the yellow dots are the host plants of the bacterium. **C.** Map of the symptomatic trees located in the 100-m area around the plantation, presenting typical *X. fastidiosa* leaf scorches. The green dots are the asymptomatic plants, the yellow dots the symptomatic plants and the red dots the symptomatic plants that are host plants of the bacterium. The maps were created with the QGIS software with maps from Google Earth, Imagery ©2018, DigitalGlobe.

Regarding the rosemary spy plants, molecular tests carried out over four years have not detected any bacteria in the collected samples. The rosemary have suffered from the heat and many of them died. In May of the first year, the 12 rosemary planted in the campus were already all desiccated. The following year they were replaced, as well as six rosemary plants located in the demarcated area. However, they did not last one year. Soil tilling performed in the demarcated area by the local gardeners also removed several plants from the ground. Only 12 out of 44 rosemary survived the four years of the experiment. The first year, symptoms already started to appear in May, and at the end of the first season, two third of the plants presented typical *Xylella*-symptoms, starting with chlorosis at the tip of the leaves, which extends to all the leaf surface and which turned necrotic (Fig. 6).

**Figure 6.**
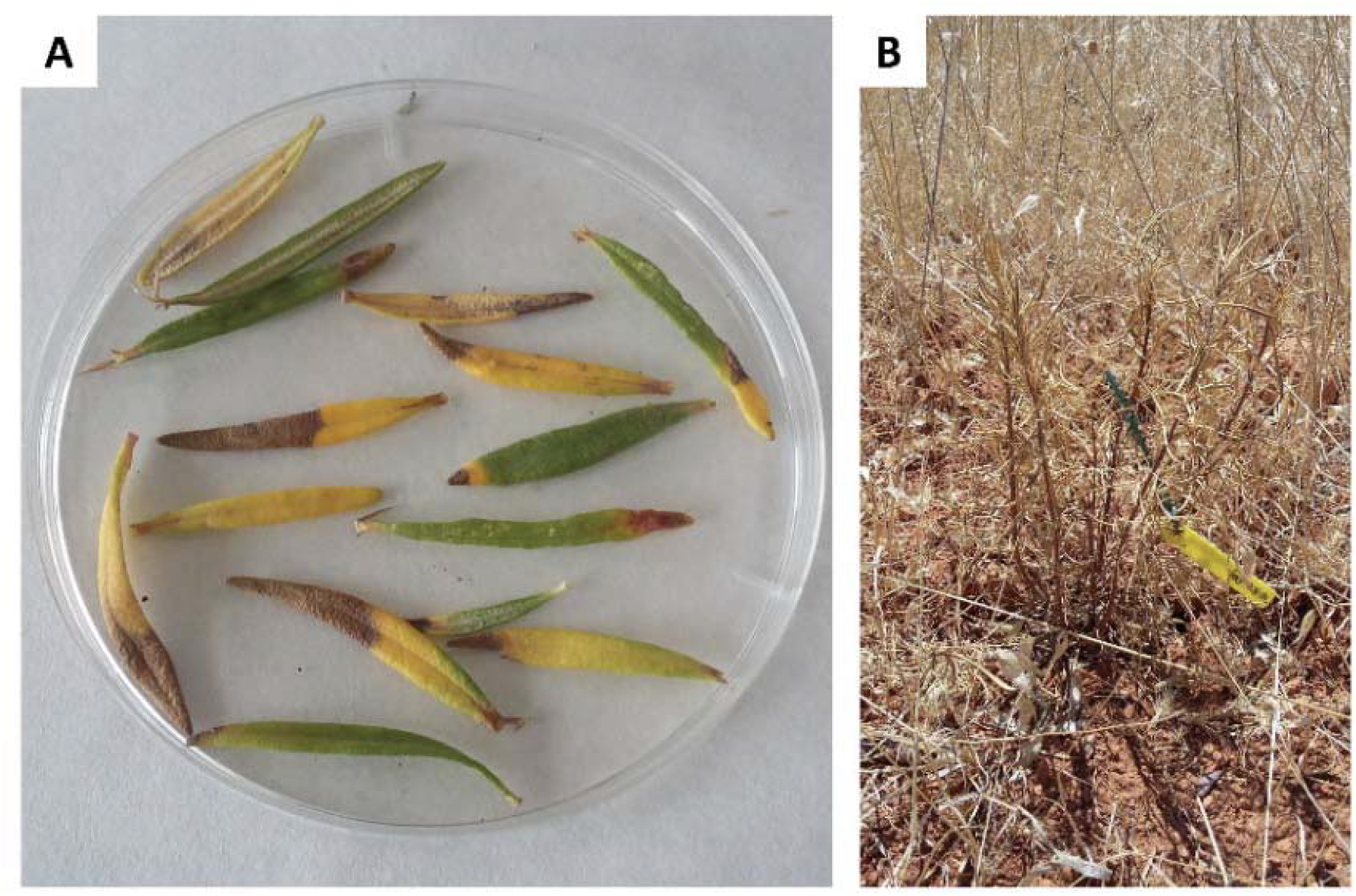
Rosemary health state. **A.** Sampled leaves of rosemary presenting *X. fastidiosa-typical* leaf scorch symptoms (May 2018). **B.** Dry and dead rosemary on the field (July 2018).

### Insect sampling

Molecular tests carried out over four years have never detected any bacteria in the collected insects of the demarcated area.

During the first season, the amount of sampled insects of both species fluctuated depending on the month. This fluctuation can be observed in Fig. 7. In March, the foam produced by the nymphs could be easily observed and in total, 40 nymphs of *P. spumarius* (1.9 nymphs/m^2^, mainly at nymphal stage 3-4) and 89 nymphs of *N. campestris* (4.2 nymphs/m^2^, mainly at nymphal stage 2-3) were sampled. The nymphs of *N. campestris* were always found on Poaceae while *P. spumarius* ones were sampled on Asteraceae (*Carduus* sp.), Euphorbiaceae and other herbaceous plants. At the beginning of May, local collaborators observed nymphs of *P. spumarius* on one *S. alba* plant in the plantation, as well as two adults of *P. spumarius* on *P. domestica*.

**Figure 7.**
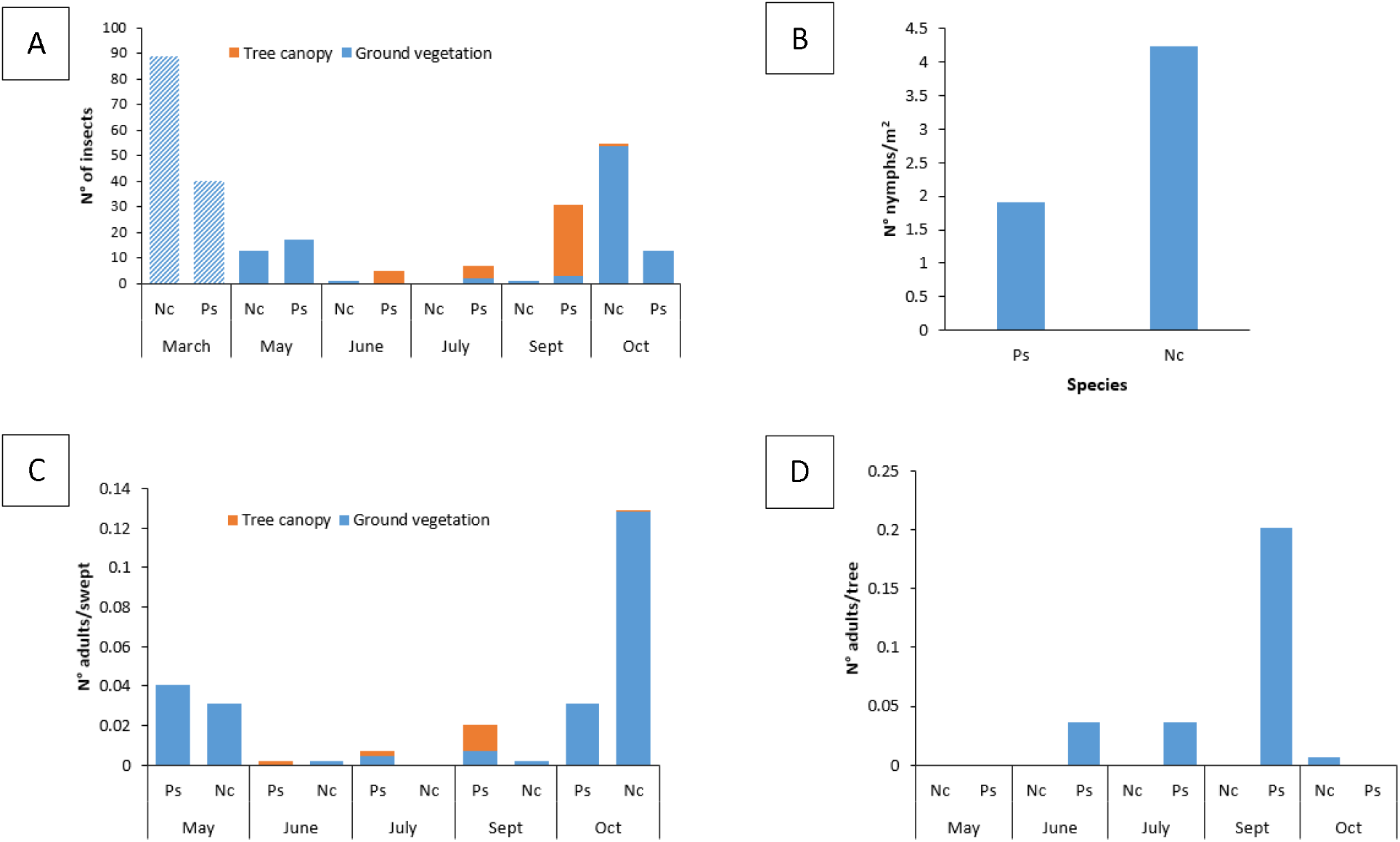
*Philaenus spumarius* (Pc) and *Neophilaenus campestris* (Nc) samples in 2018 in the 100 m-area around the sentinel plantation. **A.** Number of insects sampled through the different months. The striped pattern represents the nymphs and the plain pattern represents the adults. **B.** Number of nymphs per m sampled in March. **C.** Number of adults per swept sampled through the different months. **D.** Number of adults per tree (wild olive, almond or carob tree) sampled during the different months.

At the end of May, the adult stage was already present and the sampling on the ground vegetation revealed less individuals than when nymphs were sampled the previous months. The number of adults per swept was below one, with 0.04 *P. spumarius/*swept and 0.03 *N. campestris*/swept. In June, the herbaceous layer had dried and almost no insects were found in the ground vegetation. Very few insects were also sampled in the tree canopy. In September, more *P. spumarius* adults were sampled in the tree canopy, however the number remained low with about 0.2 adultsltree. In October, new fresh herbs had grown and the highest number of *N. campestris* over the season was reached in the ground vegetation (0.13 adults/swept), while a similar density as the one sampled in May has been found for *P. spumarius* (0.03 adults/swept).

The following years, the amount of insects collected around the plantations varied between months and years (Fig. 8) with a maximum in October 2020 of 0.06 *P. spumarius/*swept and 0.13 *N. campestris/*swept, sampled in the ground vegetation for both species. In total, four *P. spumarius* in October 2019, one *P. spumarius* in October 2020 and one *N. campestris* in October 2020 were found in the herbaceous layer of the sentinel plantation, showing that few insects were also circulating among the trees.

**Figure 8.**
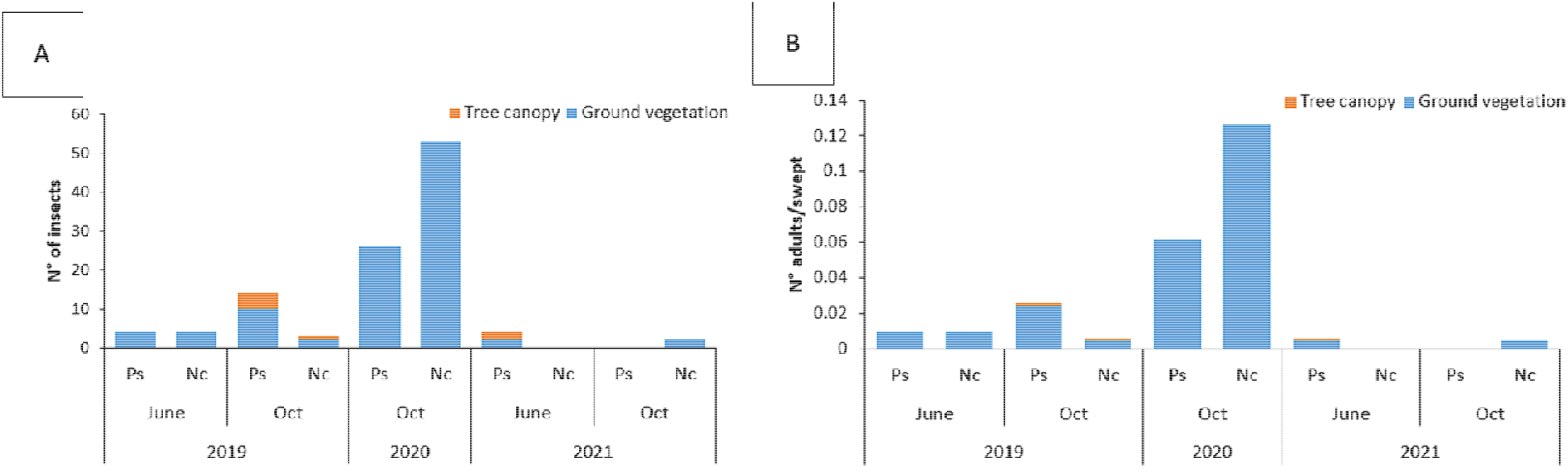
*Philaenus spumarius* (Pc) and *Neophilaenus campestris* (Nc) samples in 2019, 2020 and 2021 in the 100 m-area around the sentinel plantation. **A.** The total number of collected adult insects. **B.** Number of adults per sweep.

### The sentinel plants

Molecular tests carried out over four years have never detected any bacteria in the collected samples of the sentinel plants.

Nevertheless, first symptoms on *S. alba* already started to appear in June of the first year (2018) with some slight necrosis at the leaf margins of some of the plants. In July of that year, 78 % (21/27 plants) of the willows had slight symptoms, while in October, 96 % (26/27 plants) presented leaf necrosis starting from the tip, sometimes followed by chlorosis (Fig. 9). Regarding *P. domestica*, slight chlorosis followed by necrosis at leaf margins started to appear in July 2018 on five of the plants (Fig. 9). In October of the same year, ten plants had slight symptoms and two had moderate symptoms of chlorosis and necrosis of leaf margins. Finally, concerning *Q. petraea*, first typical necrosis on leaf margins started to appear in September of the first year. In October, these symptoms were more widespread affecting 30% of the plants (8/27 plants) and consisted in typical necrosis of leaf margins with a chlorotic halo (Fig. 9), while two plants completely died. The following years, the same symptoms started to appear on the new growing leaves, mainly on *S. alba* and *Q. petraea*. On *P. domestica*, typical leaf symptoms were less frequent; however, this species presented more defoliation. The second year, the extremity of the principal stem of five plum trees and five willows started to die; for the three species, stem sprouts started to grow on 1-2 plants per species.

**Figure 9.**
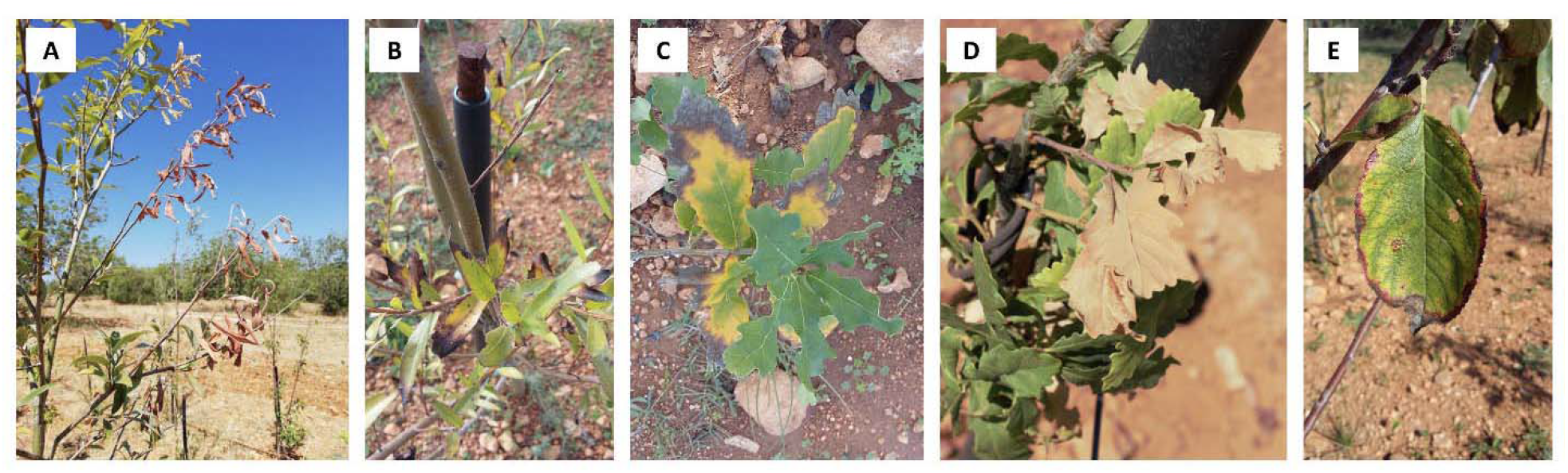
*Xylella-like* symptoms on the plants of the sentinel plantation in October of different years. **A.** On *Salix alba* in 2019. **B.** On *Salix alba* in 2018. **C.** & **D.** On *Quercus petraeae* in 2018. **E.** On *Prunus domestica* in 2018.

The summer of 2021 was declared the warmest recorded in Europe in the last 30 years, with severe heatwaves in the Mediterranean (Copernicus 2022). While the sentinel plants were already weakened by the last three hot summers despite the irrigation system, many of them died completely or partially this last year. Death was assigned after scraping the bark from several parts of the trunk. In total 14 *S. alba* plants were completely dead, and 13 had their main stem completely desiccated but had developed sprouts at the bottom that were still living. The remaining leaves showed all symptoms of necrotic and chlorotic leaf margins. Three *Q. petraea* died and almost every remaining individuals presented symptomatic leaves, while two of them had their main stem completely dead but with living sprouts. Finally, two *P. domestica* died and about twenty of them had symptomatic leaves, which consisted of leaves turning red from the margins with a degraded color, except for some leaves where the discolored margins were quite delimited. About fifteen plants had between a quarter and a half of their main stem completely dead starting from the tip. Finally, two of them had their stems completely rejected, leaving a second plant to grow from the variety Myrobolan, as the Opal variety was grafted onto this rootstock. The size measured each year was not reported here because it was biased by the death, or partial death, of the main stem.

Concerning *Q. petraea*, damage caused by the herbivores *Lachnaia septempunctata* in May-June 2018 and *Lachnaia sexpunctata* Scopoli, 1763 in June-July 2018 forced us to put their foliage under a net (Fig. 10) until mid-July to maintain them alive, but this also resulted in their inaccessibility to *X. fastidiosa* insect vectors. A pesticide (Cypermethrin 10 mL/L) also had to be applied. The following years, the situation was better and the foliage could be exposed to the environment all the season. During the monitoring, fungal-like agents were also observed on leaf surface of many individuals.

**Figure 10.**
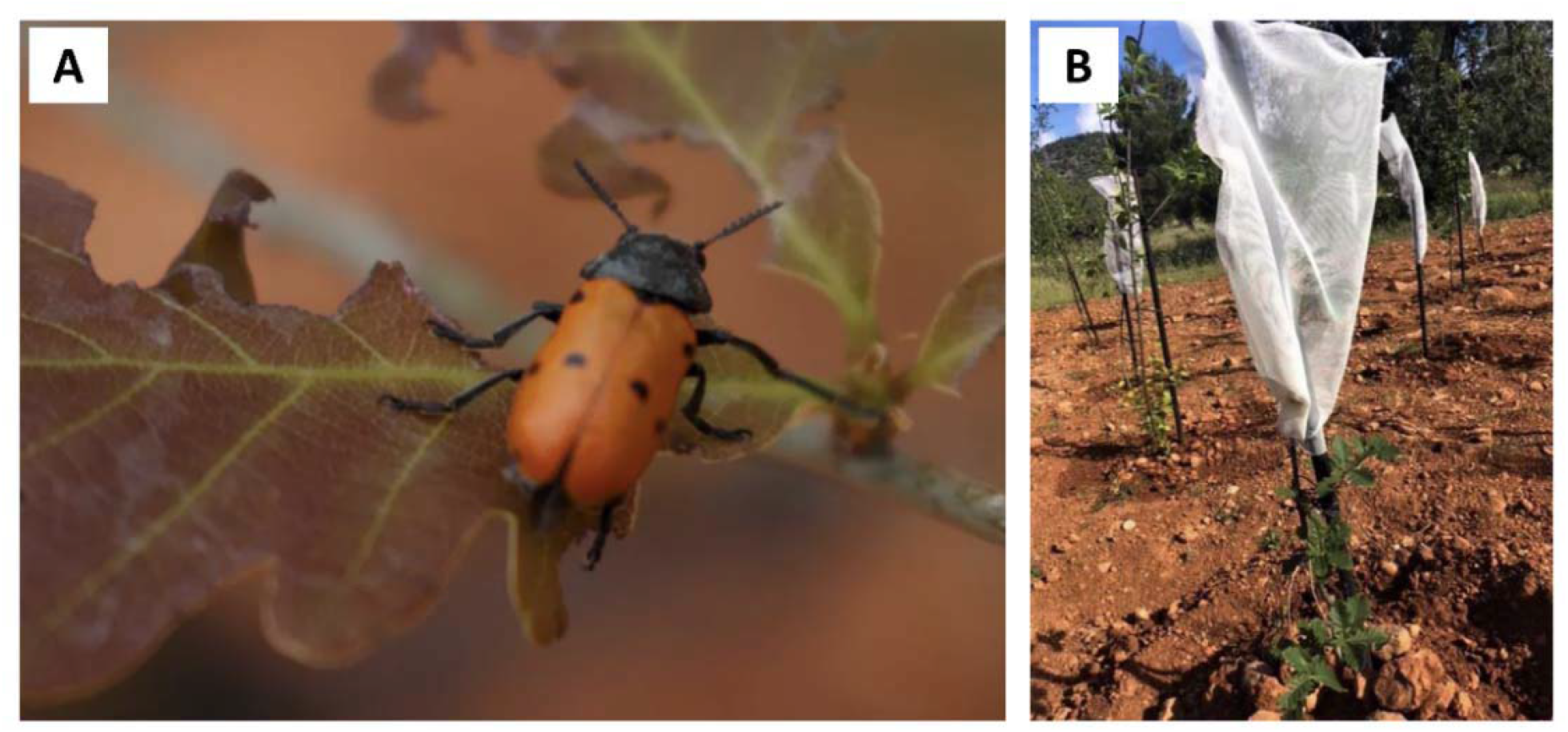
Herbivore damage on *Quercus petraea*. **A.** *Lachnaia sexpunctata* feeding on *Q. petraea* in the sentinel plantation. **B.** Net on *Q. petraea* to protect them against the herbivores.

## Discussion

During the four-year sampling and monitoring, *X. fastidiosa* was never detected in our sentinel plants nor in the collected insects. While, it is rather positive not to have any infection of this quarantine pathogen, the duration of the plantation establishment did not allow to answer the question of the potential host range of *X. fastidiosa*. In fact, besides the low infectivity pressure that had been observed on the plot, the absence of detected interaction between the bacterium and the sentinel plants does not mean that interaction could never occur (Mansfield et al. 2019) mainly given the highly specific conditions required by this plant pathogen. Instead, this study was an experimental work to learn how to combine sentinel plantation and research on *X. fastidiosa*, by exploring the constraints that were encountered to improve or redirect the method for future sentinel projects. In addition, the establishment of this plantation has provided valuable data on insect abundance and infection rates near the UIB campus, and has enabled the implementation of other parallel experiments while establishing a lasting international cooperation between the two universities.

### Complexity of sentinel plantations combined with *X. fastidiosa* research

Despite the publication of EPPO in 2020 providing guidelines for sentinel studies, only two other assays that describe themselves as sentinel plantations have been reported in the literature and both as part of the same project (Roques et al. 2015; Vettraino et al. 2015), while a third study can be characterized as one even if it does not refer as such (Rathé et al. 2014). The sentinel plantations of Roques et al. (2015) and Vettraino et al. (2015) consisted of a four-year monitoring of five European tree species, including *Quercus* spp., which had been planted in China to investigate potential new host-pest/pathogen associations that could emerge in Europe through plant trade. While the experiments allowed collecting valuable data discovering new associations, it already highlighted the complexity of the technique in terms of logistics and workload.

In our study, many constraints were faced and are reported in Table 2 with some perspectives on how the system could be improved to ease the implementation of the method. Our burdens started with permits and Italian administrations. In fact, the initial plan was to establish the plantation in the Apulian area where the first epidemic was declared. *Ex-patria* sentinel plantation studies require the movement and planting of non-native plants and they are therefore subjected to the host country’s legislative and administrative procedures for importation and planting (EPPO 2020). After more than one year of back and forth e-mails to get approval from the Italian authorities, our request was transferred to our first correspondent. Therefore, the location of the plantation was changed to Majorca, where a good collaboration with UIB allowed us to obtain the agreement of the local authorities and the university, where the plantation was to be established, in about a month. A comparative view of the full procedural pathway between our first attempt in Apulia and Majorca can be viewed in supplementary material (see supplementary file 1). Also for administrative reasons, Roques et al. (2015) were unable to establish their plot in the initially optimal climatic zone they wanted to. In their study, many plants were lost due to the delays in Chinese authorizations and imposed quarantine measures. Due to its common external border, plants with a European passport can circulate in Europe without restrictions and sentinel plantation *intra*-Europe should therefore be easier to implement (Vettraino et al. 2020). Furthermore, Vettraino et al. (2020) classified Europe as having low bureaucratic complexity concerning sentinel plantations compared to other non-European countries in a ranking they established according to the country’s bureaucratic procedures. Surprisingly, Italy was considered the least complex European responding country, in contrast to what was experienced here. However, the current sensitive issue of *X. fastidiosa* in Italy has certainly not helped to speed up the procedures. On the other hand, the government of the Balearic Islands immediately accepted our request under certain conditions, which were the compliance with the norms in force in the territory regarding *X. fastidiosa* and the prohibition of planting *Polygala myrtifolia*, initially chosen as spy plant for its high susceptibility to the bacterium. Vettraino et al. (2020) reported that most of the countries have restrictions on the import of certain plant species or genera, e.g. Roques et al. (2015) were prohibited from planting *Pinus* spp. for their sentinel plantations in China. Finally, it is worth noting that we were not able to import plants collected in semi-natural environments, such as cuttings of *S. alba*, because of the difficulty of obtaining phytosanitary passport for this type of material and all imported plants had to be purchased from Belgian nurseries in order to be certified.

**Table 2.** Constraints and perspective of using sentinel plantation for *Xylella*-research. Constraints encountered in establishing a sentinel plantation in the case of a *Xylella fastidiosa* survey and perspectives for improving the implementation of the method.

| Constraints |  | Perspectives |  |
| --- | --- | --- | --- |
| **Administrations** |  |  |  |
| - | Administrative procedures: Apulia vs. Majorca.<br>Probably impacted by the sensitive issue of *X. fastidiosa*. | - | Need for strong, organized and well-informed partnership.<br>Despite EPPO guidelines (2020), need for more homogenization of admin. procedures and interpretation of the regulations at European level (and at global level through other intercontinental organization/conventions), requiring to improve consideration towards sentinel plantations by increasing awareness of their usefulness. |
| **Legal frame of importing exotic plant material** |  |  |  |
| - | Complexity of obtaining a European passport for material collected in semi-natural environments. | - | Need to simplify the procedures at national level for obtaining passports for scientific research purposes, under verification conditions of the plant material innocuity. |
| **Legal frame of *X. fastidiosa*, as a quarantine agent** |  |  |  |
| - | No movement of plant material from infected zones. | - | Need for a proper bio-molecular processing infrastructure on site. |
| - | Removal of infected plants and vector control, decreasing infection pressure around the plantation. | - | Choosing a containment site, and not an eradication site.<br>Need for further PRA exploration if special permits could be obtain for not uprooting infected flora for scientist research purpose or for maintaining plants under certain conditions, e.g. by placing an insect-proof net on the plants to prevent spread by vectors. |
| - | Routing tests of local host plants not advised. | - | Plantation of own susceptible spy plant network. |
| - | Restrictions of planting specific plant species (*Polygala myrtifolia*). | - | / (or obtaining special permits for research purpose after PRA exploration. Need to verify plant innocuity and to sample them regularly to remove them as soon as possible in case of infection to prevent participating to the spread of the disease locally). |
| **Complexity to detect *X. fastidiosa*** |  |  |  |
| - | Concentration below threshold of methods. | - | Need to use several specific and sensitive detection methods (quantitative PCR, nested PCR...). |
| - | Irregular distribution in plants and asymptomatic plants. | - | Multiply sampling from all sides of the plant (leaves and twigs). If symptoms, prioritizing sampling of symptomatic parts. |
| - | Symptoms easily confused with ones due to other causes. | - | Validation of bacterial presence only if detected with two different methods. |
| **Length of establishment** |  |  |  |
| - | Incubation period and length of establishment of *X. fastidiosa*. | - | Long-term international financial and workload support.<br>Considering arboreta and botanical gardens studies. |
| - | Plants submitted to the unpredictability of natural conditions, with high dependence on abundance, host preference and prevalence of *X. fastidiosa* insect vectors. | - | Knowing the epidemiology of the exact sentinel location and choose a plot with high infective pressure (insect prevalence on site is measured on adults).<br>Considering targeted transmission experiments on sentinel plants with wild insects collected from naturally infected areas. |
| **Pathosystem polymorphism** |  |  |  |
| - | Investigation only of potential pathosystems based on local components.<br>For Majorca, strains: ST1, ST81, ST7; insect vectors: *P. spumarius* or *N. philaenus*; local environmental conditions. | - | Choosing location according to the strains one wants to test (extrapolating on current knowledge on which bacterial subspecies affect which plant genus can help but it is not always accurate).<br>If no preferential subspecies, choosing location with the most strains present, or multiply experiments to several areas.<br>Choosing location with the closest conditions to country of origin (environmental or insect vector population type). |
| **Abiotic and biotic stress for plants** |  |  |  |
| - | Other symptoms masking those of interest | - | Irrigation system, eventual fertilizer application. |
| - | Plant mortality limiting the experiment | - | Fitting environmental conditions of native area if possible (in case of northern European countries more complicate with *X. fastidiosa* only occurring in southern Europe, thus, considering arboreta and botanical garden studies or targeted transmission experiments in infected areas). |

The second challenge of this plantation was to keep the plants alive. The fact they were grown in an environment with different conditions including temperature and soil, brought different biotic and abiotic stresses. The life of these plants depended once again on the good cooperation on site. For example, the delay in the irrigation system establishment in the first year led the local staff to water the plants by hand every two days, carrying more than 80 L of water in cans to the plantation. Furthermore, if they had not placed mesh covering the foliage for herbivores such as *L. septempuctata* that devoured the oak leaves, the plants would have died during the first year. However, despite constant monitoring by local collaborators, plant mortality increased from year to year and stress often led, especially in willows, to a death of the main stem and the growth of new shoots at the bottom of the plant. This may have an impact on the outcome of the experiment, as the death of the potentially contaminated plant parts would lead to the death of the bacteria itself.

Here, the hurdles faced in sentinel plantation assays were coupled with the difficulties often encountered in *X. fastidiosa* studies. In fact, this bacterium is known to be fastidious for research including in its detection (Wells et al. 1987). Its concentration in plants and insects could be below the detection threshold of the different methods (Cruaud et al. 2018; EPPO 2019) and it is irregularly distributed in plants so may be missed during sampling, especially in asymptomatic plants (EFSA PLH Panel 2015; EPPO 2019). On the other hand, symptoms are not always reliable as they can easily be confused with symptoms triggered by other factors such as drought (EFSA PLH Panel 2015). Therefore, it is more than likely that other causes, such as drought or soil stress, were responsible for the typical chlorosis and necrosis of the leaf margins observed on all three species in this study, especially for such plants used to colder temperature and more humid soil, even with the irrigation provided. While an undetectable low bacterial concentration can be questioned, several studies reported that high symptomatic responses were correlated with high bacterial loads (Holland et al. 2014; Saponari et al. 2017) suggesting a greater probability of detection if symptoms were due to *X. fastidiosa* infection.

Another parameter to consider when studying host susceptibility of *X. fastidiosa* is that the incubation period can be measured over months and years (EFSA PLH Panel 2019b), indicating that time is a key element. For example, the survival time of Majorcan almond trees from bacterial infection to tree decline has been estimated around 14 years (Olmo et al. 2021). Sentinel plantation studies are already by themselves long-term assays, and superimposing the potential time required for infection of the bacteria gives us an idea of how long it takes to conduct this type of experiment. However longer incubation period does not necessarily mean lower susceptibility to the bacterium itself, since many external factors can influence it, for example the vector population. In fact, as *X. fastisiosa* is an insect vector-borne pathogen, its circulation and infection will depend from the abundance, host preference and prevalence of its insect vectors, which is adding complexity to the system compared to other sentinel studies that would for example measure the direct impact of herbivores on leaves. Moreover, a particularity of diseases caused by *X. fastidiosa* is the polymorphism of the pathosystems. In fact, different strains and bacterial subspecies will act differently with the various xylem-feeding insect species and the different host species or cultivars, leading to very specific epidemics around the world (Pierce’s disease, Citrus variegated chlorosis, Olive quick decline…) to almost no symptoms or to an endophytic presence. While the choice of the region in relation to the strains one wants to study is essential, this means that an absence or an endophytic interaction does not mean that other strains cannot be aggressive on the same plant species and cultivar. This means that there will only be an answer for a potential pathosystem related to the chosen region, but there are multitudes of other possibilities. The identified pathosystem will keep the adjective “potential” until the disease is not actually observed in the country of origin, as local environmental conditions or the presence of an effective vector will also have an impact.

A final element to be taken into account in the case of sentinel plantations with *X. fastidiosa* is the European regulation as a quarantine agent (Council Directive 2000/29; EU 2000) and the European containment and eradication measures imposed in case of detection (Regulation EU 2020/1201, EU 2020) with the establishment of a demarcated area delimiting an infected zone of at least 50 m and a buffer zone varying in terms of kilometers depending on the situation. In the infected zone, eradication measures have to be undertaken consisting of the removal of all specified host plants of *X. fastidiosa*. However, in areas in which the bacterium is considered widely established including Apulia, Corsica and Balearic Islands, lighter containment measures may be implemented as eradication is no longer considered feasible. Nevertheless, these measures still imply the removal of all the infected plants in the 50 m-zone, and an intensive surveillance within an area of at least 5 km-radius together with vector control. These measures mean that even in containment zones such as the Balearic Islands, the detection of an infected plant in sentinel plantation would lead to a control of vector population in the area, and to a decrease in the infection pressure around other plants of the plantation. Similarly, if the tested positive plant has to be removed immediately, the observation of symptom evolution and thus, the assessment of susceptibility is compromised, unless exceptional permits for scientific research are obtained. In this study, the problem did not arise because all plants tested negative. Nevertheless, we were still impacted by the consequences of the European legislation as under the containment scenario in the Balearics, local government and UIB authorities did not advise systematic test of the host plants on the campus. In fact, a positive detection would have led to the uprooting of the campus vegetation, including, as mentioned before, our plantation if special permits were not issued. These measures are considered highly severe for an area where the bacterium is widespread and separated from other regions by the sea (Olmo et al. 2021). In areas infected by *X. fastidiosa*, the possibility of not having to remove infected plants in the field for scientific research purposes deserves further exploration in terms of PRA and bureaucratic procedure. Finally, for biosafety reasons related to quarantine organisms, plant samples cannot be moved and have to be processed on site, which again requires a good logistic, local collaboration and proper infrastructure.

### Necessity of knowing the epidemiology of the exact sentinel location

The implementation of a sentinel plantation when studying a specific pest or pathogen requires knowing well the epidemiology of the exact spot of the establishment, as local environmental components have a great impact on the outcome of the experiment (Kenis et al. 2018). The location chosen for this study was probably not optimal, as it was later evidenced that *X. fastidiosa* infection pressure was low, and thus, this certainly constitutes the main reason for the lack of positive detections in insects, spy and sentinel plants in the plot. When the plantation was established on the UIB campus, the prevalence and the epidemiology of the outbreak on the island were not yet well known, which is still the case in several regions where *X. fastidiosa* has recently been detected. Positive detections were reported on the campus about hundred meters from the plantation on one *R. officinalis* plant and two olive trees (M. A. Miranda personal communication) and the health state of host plants including declining almond trees, one of the main crop affected by *X. fastidiosa* on the island, led us to suspect that the place was infected. However, due to the lack of systematic sampling after the declaration of the contention scenario in the Balearics, the presence of the bacterium could not be confirmed by testing. In addition, the quantity of nymphs sampled when choosing the location was 1.9 nymphs/m^2^ for *P. spumarius* and 4.2 nymphs/m^2^ for *N. campestris* in March, which is actually higher than the mean observed in the ground vegetation sampled through the island. López-Mercadal et al. (2021) reported an average of about 0.22 nymphs/m^2^ for *P. spumarius* in the peak of March and 0.005 nymphs/m^2^ for *N. campestris* with differences between plots and years. In our plot, the prevalence of these nymphs was null. However, this information was not relevant as the infectivity is lost with each molt (Purcell and Finlay 1979) and prevalence therefore has to be measured on adult insects to have robust data.

After deepest outbreak investigations, it appeared that the east side of the island towards Manacor was probably the most infected part. In fact, Gutiérrez Hernández and García (2018) mapped the positive records of *X. fastidiosa* detected in the Balearics by the Plant Health Section of the Department of Environment, Agriculture and Fisheries of the Government of the Balearic Islands, and showed that most of the positive samples were concentrated on the east side (Manacor, Sant Llorenç des Cardassar and Son Servera; Fig. 11) with the highest densities in agricultural and residential areas close to the main communication routes. They stressed, however, than the conducted sampling strategy could have biased this distribution, for instance, because the samples could have been collected preferentially in these more accessible areas. Based on direct field observations and using Google street view, Moralejo et al. (2020) also mapped the distribution of *Xylella*-symptomatic almond orchards and their mortality across the island, tracking their evolution since 2012 (Fig. 11). They showed a gradient from east to west, showing a moderate incidence on the site of the plantation. However, molecular testing of infected almond trees did not reveal a clear spatial pattern (Moralejo et al. 2020). In addition, highly variable incidence was encountered in different orchards (Olmo et al. 2021), hence the need of knowing the incidence and prevalence of vectors at the precise location of a sentinel plantation.

The density and prevalence of insect vectors are one of the drivers of *X. fastidiosa* infection and impact the temporal dynamics of symptom appearance (EFSA PLH Panel 2019b), as multiple and independent infections could lead to an injection of a higher bacterial load and a decrease in the incubation period (Daugherty and Almeida 2009). The damage in the Balearics are the consequence of almost 20 years of infection (Moralejo et al. 2020), suggesting that the infection pressure could be too low to conduct sentinel plantation experiments. In fact, the abundance of nymphs and sampled adults as well as the prevalence of insects are lower than the values encountered in the infected areas of Apulia where the outbreak was more drastic. A prevalence of 23% was reported in Majorca (López-Mercadal et al. 2021) compared to up to 71% detected in an Apulian olive grove (Cornara et al. 2016a). Similarly, higher densities of vectors were measured in Apulia with 7 to 39 nymphs of *P. spumarius*/m^2^ in olive orchards (Bodino et al. 2019), about 7 adults/olive trees and 0.5 adults/swept in weeds recorded during the respective seasonal peaks (Cornara et al. 2016b), however, with heterogeneity identified among the orchards studied (Bodino et al. 2019). In our plot, the adult density varied according to the seasonal estivation and ground drying pattern of Mediterranean regions (Cornara et al. 2016b; López-Mercadal et al. 2021). It barely reached a maximum of 0.04 *P. spumarius*/swept in May and 0.2 *P. spumarius*/tree in September 2018, while the average reported through the island was below 0.1 adults/swept in ground cover, tree canopy and border vegetation (López-Mercadal et al. 2021). In addition, *N. campestris* was not considered as a significant vector due to its very low presence on the tree canopy (Lopez-Mercadal et al. 2021). Moreover, the soil around the plantation was plowed almost every year, as common management on the island, which besides destroying several rosemary spy plants, probably decreased insect movement around the plot even with sowing of ground vegetation the second year. In fact, tillage is a technique of vector control reducing the number of vector/m^2^ (Bodino et al. 2019; EFSA PLH Panel 2019b). In the study of Kenis et al. (2018), their plantation located at the edge of the forest took less time to be infested than another one situated in an agricultural-peri-urban area, highlighting again the impact of high local circulation of pests and pathogens on the time and outcome of the assay.

### Sentinel plantations as an efficient tool for *X. fastidiosa* research in specific situations

Even in locations with high infection pressure, the efficiency of the sentinel plantation in the case of *X. fastidiosa* host range investigation is questioned due to the ratio results/time-workload. Yet the sentinel plantation method is currently being used in Apulia for the screening of olive cultivars coming from various Mediterranean olive-growing areas (Spain, Tunisia, Greece, etc.) by exposing them to the natural pressure of inoculum in heavily infected field (XF-ACTORS 2017; Saponari, et al. 2019). The previous finding of the mild symptoms on the Leccino and FS17^®^ olive cultivars adjacent to severely affected orchards, motivated the study (Boscia et al. 2017). Approximately 100 different genotypes were planted and are currently under evaluation in different plots, actually making the Apulian region home to one of the largest sentinel plantation of all time. This study is promising and is considered necessary for long-term management of *X. fastidiosa* in olive growing regions as preliminary data show already differences in susceptibility in various cultivars (EFSA PLH Panel 2019b; Saponari et al. 2019). However, it highlights the long-term commitment required as the survey started in 2015 and is still ongoing. The project is part of a research program funded by the European Union’s Horizon 2020 Research and Innovation Program, which explains how a project of this magnitude could be established and which underlines the need for long-term consistent international support for the implementation of such experiments. The success of this plantation, in addition to the selection of highly infected plots, also comes from the fact that the tested potential hosts are related to the STs present in the environment. The Apulian ST53 being highly aggressive on olives, it is obvious to carry out olive plant susceptibility in this area. However, other *X. fastidiosa* infected regions as Balearic Islands and Corsica could be interesting to study the susceptibility to other STs, as three STs belonging to two subspecies coexist in Majorca while only one in the Apulian region.

Thus, the Apulia study proved the usefulness of sentinel plantations in the context of *X. fastidiosa*. However, it would be less relevant to conduct these studies in certain situations. There should be, for example, similarities between the climatic conditions of the two regions involved in the sentinel studies to minimize the impact of external factors. So far, the bacterium has only been found established in southern Europe, in regions with a Mediterranean type of climate, and these studies would therefore be less suitable for northern European countries, as differences in environmental conditions could lead to weakening or even death of the plants and to misidentification of the cause of potential symptoms. Nevertheless, this tool remains very valuable and should be considered for studies on *X. fastidiosa*, as other techniques for screening potential hosts of this pathogen are also discussed. Among these techniques, mechanical inoculation shows a low rate of success, even in susceptible hosts (Prado et al. 2008; EFSA PLH Panel 2019b) as this method artificially reproduce infection while in the environment, only xylem-specialized insect vectors have the capacity to infect plants (Almeida et al. 2005). Working with insect vectors is therefore a more relevant way of conducting experiments. However, besides the biosafety risk it could represent for *Xylella*-free regions and the need for proper infrastructure, the very act of infecting an insect is a challenge. Other experiments consisting in grafting more than 400 olive genotypes on infected trees were conducted in parallel of sentinel plantation in Apulia to short incubation period and time imposed by insect traits (Saponari et al. 2019). However, in addition to also being an artificial way of infection, it requires the availability of appropriate infected graft material. Therefore, sentinel plantation has its advantages and has to be considered a valuable complementary tool in certain situations.

In these situations, this study has provided a complete methodology to monitor the bacterium circulation through the sentinel plants. The use of spy plants is certainly useful if sampling of susceptible vegetation is not possible in the nearby area. In other cases, sampling of local flora may be sufficient, although it does not ensure real-time circulation of the bacteria, as the current state of the local flora could be the result of infection from the past (Moralejo et al. 2020). The use of small perennial plants may facilitate sampling, as bacteria are distributed irregularly in the plant. The species or mix of species must be adapted to local conditions, susceptible to the bacterial strains being investigated and favored by local vectors. In this study, *R. officinalis* was chosen as it was reported infected with the European STs of subsp. *multiplex* and subsp. *pauca* (ST6, ST7, ST53, ST80, ST81 and ST87) and was found infected in Majorca with the ST81 (EFSA 2022). In addition, in America, the bacterium was detected on this plant species close to *X. fastidiosa* subsp. *fastidiosa*-infected vines (Freitag 1951).

### Conducting sentinel studies differently to assess host range up North

Sentinel studies can also be carried out differently to study host range in countries that cannot match closely the environmental conditions of the potential location. First, arboreta and botanical gardens are still an option of studying exotic host range in naturally infected environments. However, as a detectable infection depends on the density and prevalence of *X. fastidiosa* insect vectors (Daugherty and Almeida 2009), the use of this method could also be discussed as these areas are often subjected to phytosanitary management. One advantage of these studies regarding *X. fastidiosa* would be that plants are grown in these sites for a long time, increasing the success concerning potential latent periods or low bacterial load potentially enabling detection. In addition, the study of Groenteman et al. (2015) has shown promising results for *X. fastidiosa* research by sampling in botanical gardens. They managed to discover 28 New Zealander plant species infected by *X. fastidiosa*, including several visited by the insect vector *Homalodisca vitripennis* Germar, 1821, in Californian botanical gardens where the disease is well established. They also found parasites capable of controlling the vector on these plant species in the aim of a biocontrol early-response strategy in case *H. vitripennis* invade New-Zealand.

A second way would be to carry out transmission experiments with naturally infected vectors in contaminated regions to bypass the problems of biosecurity imposed by *Xylella*-free areas and the difficulty of infecting insects. Compared to standard sentinel plantations, these experiments allow to reduce the dependence on vector density and on insect feeding preferences. In fact, although *P. spumarius* is considered a polyphagous species and was observed feeding on the three studied sentinel plants in their area of distribution, it is possible that in the sentinel country, these insects are more interested in native vegetation. Native plants could therefore compete with the exotic sentinel ones, potentially resulting in fewer vector feeding events decreasing the bacterial transmission probability. Even if vector preferences are biased and that natural conditions are therefore not fully met, these experiments can still be considered as sentinel studies since they consist in *ex-patria* plants sent to study the impact of exotic organisms in areas in which they occur. This has been done in Majorca as a complementary experiment where 20 new cuttings of *S. alba* and of *P. tremula* have been sent from Belgium to the UIB campus (Casarin et al. submitted). There, transmission experiments with naturally infected *P. spumarius* were conducted in an insect-proof greenhouse and revealed positive infection on *S. alba*, proving the higher efficiency of the technique compared to sentinel plantation.

Finally, sentinel plantings “*in-patria*” (Eschen et al. 2019; or “sentinel nurseries”, sensu Vettraino et al. 2017) consist in planting native traded plants without phytosanitary treatments on its own land to monitor pests and pathogens which could be spread through international trade (Vettraino et al. 2017). They obviously do not have the same objective as *ex-patria* plantation that informs PRA of organisms that are not yet present in a given area. Rather, they consist of surveillance for a known pathogen for which possible entry and dispersal pathways have been identified (Mansfield et al. 2019) and they still represent valuable sentinel assays to be conducted in the aim of early detection of *X. fastidiosa* in new regions. The major difference with a standard commercial nursery is that no pest control measures are implemented on these plants (EPPO 2020) so that it is possible for the vectors to reach the plants and for the plants to get infected if *X. fastidiosa* is introduced in the area. For this strategy to be effective, these plantations have to be established in strategic locations where the bacterium is the most likely to enter. The “plant for planting” pathway being the main entrance for exotic organisms including *X. fastidiosa* (Liebhold et al. 2012; EFSA PLH Panel 2018), their locations in/close to nurseries or other plant commercial places would be relevant. In addition, these plantations must consist of known host plants that have a high probability to be the first infected when the bacterium enters an area and if possible, to be highly susceptible for the infection to be visible and easily detectable. For example, the Auckland Botanic Garden had set up a sentinel plot of myrtle plants to detect the potential arrival of the myrtle rust (*Puccinia psidii* Winter, 1884) as early as possible in New Zealand, as the fungus was prevalent in Australia at the time (Barham et al. 2015). Similarly, one can imagine planting a network of *P. myrtifolia* near nurseries, previously tested for innocuity, which are regularly monitored for potential contamination by *X. fastidiosa*. Obviously, these susceptible plants should be tested carefully and regularly to provide the benefits of early detection while preventing them from serving as inoculum for disease establishment (Mansfield et al. 2019).

## Conclusion

In conclusion, this study is an experimental work highlighting that sentinel plantations are not easy to implement in the case of *X. fastidiosa*, but that they are complementary to other studies and that they could provide valuable information on host interactions when some conditions are met. This work proposes a methodology to monitor future sentinel plantations and it suggests other ways of conducting sentinel experiments for screening host range or for early detection of *X. fastidiosa* in new areas.

## Acknowledgments

We would like to acknowledge all the people who helped in the establishment of the sentinel plantation and the irrigation system, and who contributed to maintain the plants alive, especially the UIB gardeners and technicians, Amélie Emond, Maria Antònia Tugores, Pau Mercadal, Carlos Barceló and Noelia Barros. We also want to thank all the people who irrigated manually with bravery the plantation during the first year. We are thankful to Sofia Delgado and Claudia Paredes for their laboratory support. Finally, we are grateful to UIB authorities and to the government of the Balearic Islands for the authorization to carry out these assays.

## Author contributions

All authors designed the sentinel plantation assay. JLM and MAA took care of the plantation during the four years. NC and SH carried out the detection tests. The first author wrote the first draft of the manuscript and the last author provided the comparative view of administrative procedures in supplementary file 1. All authors commented, improved previous versions of the manuscript, and read and approved the final version.

## References

Akbulut S and Stamps WT (2012) Insect vectors of the pinewood nematode: a review of the biology and ecology of *Monochamus* species. Forest Pathology, 42(2), 89–99. https://doi.org/10.1111/J.1439-0329.2011.00733.X

Almeida RPP, Blua, MJ, Lopes, JRS, Purcell AH (2005) Vector transmission of *Xylella fastidiosa*: Applying fundamental knowledge to generate disease management strategies. Annals of the Entomological Society of America, 98(6), 775–786. https://doi.org/10.1603/0013-8746(2005)098[0775:VTOXFA]2.0.CO;2

Aukema JE, Leung B, Kovacs K, Chivers C, Britton KO (2011) Economic impacts of non-native forest insects in the continental United States. PLoS ONE, 6(9), 24587. https://doi.org/10.1371/journal.pone.0024587

Barham E, Sharrock S, Lane C, Baker R (2016) The International Plant Sentinel Network: A tool for regional and national plant protection organizations. EPPO Bulletin, 46(1), 156–162. https://doi.org/10.1111/epp.12283

Blossey B and Notzold R (1995) Evolution of increased competitive ability in invasive nonindigenous plants: A hypothesis. The Journal of Ecology, 83(5), 887. https://doi.org/10.2307/2261425

Bodino N, Cavalieri V, Dongiovanni C, Plazio E, Saladini MA, Volani S, Simonetto A, Fumarola G, Di Carolo M, Porcelli, F, Gilioli G, Bosco D (2019) Phenology, seasonal abundance and stage-structure of spittlebug (Hemiptera: Aphrophoridae) populations in olive groves in Italy. Scientific Reports 2019 9:1, 9(1), 1–17. https://doi.org/10.1038/s41598-019-54279-8

Boscia D, Altamura G, Ciniero A, Di Carolo M, Dongiovanni C, Fumarola G, Giampetruzzi A, Greco P, La Notte P, Loconsole G, Manni F, Melcarne G, Montilon V, Morelli M, Murrone N, Palmisano F, Pollastro P, Potere O, Roseti V, … Martelli GP (2017) Resistenza a *Xylella fastidiosa* in diverse cultivar di olivo. L’ Informatore Agrario, 11. https://doi.org/10.5281/ZENODO.495708

Bosso L, Di Febbraro M, Cristinzio G, Zoina A, Russo D (2016) Shedding light on the effects of climate change on the potential distribution of *Xylella fastidiosa* in the Mediterranean basin. Biological Invasions, 18(6), 1759–1768. https://doi.org/10.1007/S10530-016-1118-1/FIGURES/4

Brasier CM (2008) The biosecurity threat to the UK and global environment from international trade in plants. Plant Pathology, 57(5), 792–808. https://doi.org/10.1111/J.1365-3059.2008.01886.X

Brasier CM and Buck KW (2001) Rapid evolutionary changes in a globally invading fungal pathogen (Dutch elm disease). Biological Invasions, 3(3), 223–233. https://doi.org/10.1023/A:1015248819864924

Britton KO, White P, Kramer A, Hudler G (2010) A new approach to stopping the spread of invasive insects and pathogens: early detection and rapid response via a global network of sentinel plantings. New Zealand Journal of Forestry Science, 40, 109–114. http://www.scopus.com/inward/record.url?eid=2-s2.0-77955440077&partnerID=tZOtx3y1

Chatterjee S, Almeida RPP and Lindow S (2008) Living in two worlds: The plant and insect lifestyles of *Xylella fastidiosa*. Annual Review of Phytopathology, 46(1), 243–271. https://doi.org/10.1146/annurev.phyto.45.062806.094342

Colautti RI, Ricciardi A, Grigorovich IA, MacIsaac HJ (2004) Is invasion success explained by the enemy release hypothesis? Ecology Letters, 7(8), 721–733. https://doi.org/10.1111/J.1461-0248.2004.00616.X

Copernicus (2022) Copernicus: Globally, the seven hottest years on record were the last seven; carbon dioxide and methane concentrations continue to rise | Copernicus. https://climate.copernicus.eu/copernicus-globally-seven-hottest-years-record-were-last-seven [Accessed on 10.06.2022]

Cornara D, Cavalieri V, Dongiovanni C, Altamura G, Palmisano F, Bosco D, Porcelli F, Almeida RPP, Saponari M (2016a) Transmission of *Xylella fastidiosa* by naturally infected *Philaenus spumarius* (Hemiptera, Aphrophoridae) to different host plants. Journal of Applied Entomology, 141(1-2), 80–87. https://doi.org/10.1111/jen.12365

Cornara D, Saponari M, Zeilinger AR, de Stradis A, Boscia D, Loconsole G, Bosco D, Martelli GP, Almeida RPP, Porcelli F (2016b) Spittlebugs as vectors of *Xylella fastidiosa* in olive orchards in Italy. Journal of Pest Science, 90(2), 521–530. https://doi.org/10.1007/S10340-016-0793-0/TABLES/3

Cruaud A, Gonzalez AA, Godefroid M, Nidelet S, Streito JC, Thuillier JM, Rossi JP, Santoni S, Rasplus JY (2018) Using insects to detect, monitor and predict the distribution of *Xylella fastidiosa:* a case study in Corsica. Scientific Reports 2018 8:1, 8(1), 1–13. https://doi.org/10.1038/s41598-018-33957-z

Cunty A, Legendre B, de Jerphanion P, Dousset C, Forveille A, Paillard S, Olivier V (2022) Update of the *Xylella fastidiosa* outbreak in France: two new variants detected and a new region affected. European Journal of Plant Pathology, 1–6. 961 https://doi.org/10.1007/S10658-022-02492-Z/FIGURES/2

Daugherty MP and Almeida RPP (2009) Estimating *Xylella fastidiosa* transmission parameters: decoupling sharpshooter number and feeding period. Entomologia Experimentalis et Applicata, 132(1), 84–92. https://doi.org/10.1111/J.1570-7458.2009.00868.X

Denancé N, Legendre B, Briand M, Olivier V, de Boisseson C, Poliakoff F, Jacques MA (2017) Several subspecies and sequence types are associated with the emergence of *Xylella fastidiosa* in natural settings in France. Plant Pathology, 66(7), 1054–1064. https://doi.org/10.1111/ppa.12695

Diagne C, Leroy B, Vaissière AC, Gozlan RE, Roiz D, Jarić I, Salles JM, Bradshaw CJA, Courchamp F (2021) High and rising economic costs of biological invasions worldwide. Nature 2021 592:7855, 592(7855), 571–576. https://doi.org/10.1038/s41586-021-03405-6

EC (European Commission) (2000) Council Directive 2000/29/EC of 8 May 2000 on protective measures against the introduction into the Community of organisms harmful to plants or plant products and against their spread within the Community. Official Journal of the European Communities, 169, 1–112. http://data.europa.eu/eli/dir/2000/29/oj

EC (European Commission) (2020) Commission Implementing Regulation (EU) 2020/1201 of 14 August 2020 as regards measures to prevent the introduction into and the spread within the Union of *Xylella fastidiosa* (Wells et al.). Official Journal of the European Union. http://data.europa.eu/eli/reg_impl/2020/1201/oj985

EFSA (European Food Safety Authority) (2021) Pest survey card on *Xylella fastidiosa*. EFSA Supporting Publications, 16(6). https://doi.org/10.2903/SP.EFSA.2019.EN-1667

EFSA (European Food Safety Authority) (2022) Update of the *Xylella* spp. host plant database - systematic literature search up to 30 June 2021. EFSA Journal, 20(1). https://doi.org/10.2903/J.EFSA.2022.7039

EFSA PLH Panel (EFSA Panel on Plant Health) (2015) Scientific Opinion on the risks to plant health posed by *Xylella fastidiosa* in the EU territory, with the identification and evaluation of risk reduction options. EFSA Journal, 13(1), 3989. https://doi.org/10.2903/j.efsa.2015.3989

EFSA PLH Panel (EFSA Panel on Plant Health) (2016) Four statements questioning the EU control strategy against *Xylella fastidiosa*. EFSA Journal, 14(3). https://doi.org/10.2903/j.efsa.2016.4450

EFSA PLH Panel (EFSA Panel on Plant Health) (2018) Updated pest categorisation of *Xylella fastidiosa*. EFSA Journal, 16(7). https://doi.org/10.2903/j.efsa.2018.5357

EFSA PLH Panel (EFSA Panel on Plant Health) (2019a) Effectiveness of in planta control measures for *Xylella fastidiosa*. EFSA Journal, 17(5), e05666. https://doi.org/10.2903/J.EFSA.2019.5666

EFSA PLH Panel (EFSA Panel on Plant Health) (2019b) Update of the Scientific Opinion on the risks to plant health posed by *Xylella fastidiosa* in the EU territory. EFSA Journal, 17(5). https://doi.org/10.2903/j.efsa.2019.5665

EPPO (European and Mediterranean Plant Protection) (2020) PM 3/91(1) Sentinel woody plants: concepts and application. EPPO Bulletin, 50(3), 429–436. https://doi.org/10.1111/EPP.12698

EPPO (European and Mediterranean Plant Protection Organization) (2019) PM 7/24 (4) *Xylella fastidiosa*. EPPO Bulletin, 49(2), 175–227. https://doi.org/10.1111/EPP.12575

Eschen R, O’Hanlon R, Santini A, Vannini A, Roques A, Kirichenko N, Kenis M (2019) Safeguarding global plant health: the rise of sentinels. Journal of Pest Science, 92(1), 29–36. https://doi.org/10.1007/s10340-018-1041-6

EUROPHYT Online (2022) The European Union Notification System for Plant Health Interceptions. https://ec.europa.eu/food/plant/plant_health_biosecurity/europhyt/interceptions_en [Accessed on 24.04.2022]

Freitag JH (1951) Host range of the Pierce’s disease virus of Grapes as determined by insect transmission. Phytopathology, 41(10). https://www.cabdirect.org/cabdirect/abstract/19521100495

Groenteman R, Forgie SA, Hoddle MS, Ward DF, Goeke DF, Anand N (2015) Assessing invasion threats: novel insect-pathogen-natural enemy associations with native New Zealand plants in southern California. Biological Invasions, 17(5), 1299–1305. https://doi.org/10.1007/s10530-014-0804-0

Gutiérrez Hernández O and García LV (2018) Incidencia de *Xylella fastidiosa* en las Islas Baleares y distribución potencial en la península ibérica. Investigaciones Geográficas (Esp), (69), 55–72. https://www.redalyc.org/articulo.oa?id=17656164004

Harper SJ, Ward LI, Clover GRG (2010) Development of LAMP and real-time PCR methods for the rapid detection of Xylella *fastidiosa* for quarantine and field applications. Phytopathology, 100(12), 1282–1288. https://doi.org/10.1094/PHYTO-06-10-0168

Holland RM, Christiano RSC, Gamliel-Atinsky E, Scherm H (2014) Distribution of *Xylella fastidiosa* in Blueberry Stem and Root Sections in Relation to Disease Severity in the Field. Plant Disease, 98(4), 443–447. https://doi.org/10.1094/PDIS-06-13-0680-RE

Keane RM and Crawley MJ (2002) Exotic plant invasions and the enemy release hypothesis. Trends in Ecology & Evolution, 17(4), 164–170. https://doi.org/10.1016/S0169-5347(02)02499-0

Kenis M, Rabitsch W, Auger-Rozenberg MA, Roques A (2007) How can alien species inventories and interception data help us prevent insect invasions? Bulletin of Entomological Research, 97(5), 489–502. https://doi.org/10.1017/S0007485307005184

Liebhold AM, Brockerhoff EG, Garrett LJ, Parke JL, Britton KO (2012) Live plant imports: the major pathway for forest insect and pathogen invasions of the US. Frontiers in Ecology and the Environment, 10(3), 135–143. https://doi.org/10.1890/110198

López-Mercadal J, Delgado S, Mercadal P, Seguí G, Busquets A, Gomila M, Lester K, Kenyon DM, Pérez R, Paredes-Esquivel C and Miranda MA (2021) Collection of data and information in Balearic Islands on biology of vectors and potential vectors of *Xylella fastidiosa* (GP/EFSA/ALPHA/017/01). EFSA Supporting Publications, 18(10), 6925E. https://doi.org/10.2903/sp.efsa.2021.EN-6925

Manfredini F, Grozinger CM, Beani L (2013) Examining the “evolution of increased competitive ability” hypothesis in response to parasites and pathogens in the invasive paper wasp *Polistes dominula*. Naturwissenschaften, 100(3), 219–228. https://doi.org/10.1007/S00114-013-1014-9/FIGURES/5

Mansfield S, Mcneill MR, Aalders LT, Bell NL, Kean JM, Barratt BIP, Boyd-Wilson K, Teulon DAJ (2019) The value of sentinel plants for risk assessment and surveillance to support biosecurity. NeoBiota, 48, 1–24. https://doi.org/10.3897/neobiota.48.34205

Meurisse N, Rassati D, Hurley BP, Brockerhoff EG, Haack RA (2019) Common pathways by which non-native forest insects move internationally and domestically. Journal of Pest Science, 92(1), 13–27. https://doi.org/10.1007/S10340-018-0990-0/FIGURES/1

Migliorini D, Ghelardini L, Tondini E, Luchi N, Santini A (2015) The potential of symptomless potted plants for carrying invasive soilborne plant pathogens. Diversity and Distributions, 21(10), 1218–1229. https://doi.org/10.1111/ddi.12347

Minsavage GV, Thompson CM, Hopkins DL, Leite RMVBC, Stall RE (1994) Development of a polymerase chain reaction protocol for detection of *Xylella fastidiosa* in plant tissue. Phytopathology 84(5), 456–461. https://doi.org/10.1094/Phyto-84-456

Moralejo E, Gomila M, Montesinos M, Borràs D, Pascual A, Nieto A, Adrover F, Gost PA, Seguí G, Busquets A, Jurado-Rivera JA, Quetglas B, García J de D, Beidas O, Juan A, Velasco-Amo MP, Landa BB, Olmo D (2020) Phylogenetic inference enables reconstruction of a long-overlooked outbreak of almond leaf scorch disease (*Xylella fastidiosa*) in Europe. Communications Biology 2020 3:1, 3(1), 1–13. https://doi.org/10.1038/s42003-020-01284-7

Nunney L, Ortiz B, Russell SA, Sánchez R, Stouthamer R (2014) The complex biogeography of the plant pathogen *Xylella fastidiosa*: Genetic evidence of introductions and subspecific introgression in Central America. PLoS ONE, 9(11), 112463. https://doi.org/10.1371/journal.pone.0112463

Olmo D, Nieto A, Adrover F, Urbano A, Beidas O, Juan A, Marco-Noales E, López MM, Navarro I, Monterde A, Montes-Borrego M, Navas-Cortés 1102 JA, Landa BB (2017) First detection of *Xylella fastidiosa* infecting cherry (*Prunus avium*) and *Polygala myrtifolia* plants in Mallorca Island Spain. Plant Disease, 101(10), 1820. https://doi.org/10.1094/PDIS-04-17-0590-PDN

Olmo D, Nieto A, Borràs D, Montesinos M, Adrover F, Pascual A, Gost PA, Quetglas B, Urbano A, García J de D, Velasco-Amo MP, Olivares-García C, Beidas O, Juan A, Marco-Noales E, Gomila M, Rita J, Moralejo E, Landa BB (2021) Landscape Epidemiology of *Xylella fastidiosa* in the Balearic Islands. Agronomy 2021, Vol. 11, Page 473, 11(3), 473. https://doi.org/10.3390/AGRONOMY11030473

Parker IM, Simberloff D, Lonsdale WM, Goodell K, Wonham M, Kareiva PM, Williamson MH, von Holle B, Moyle PB, Byers JE, Goldwasser L (1999) Impact: toward a framework for understanding the ecological effects of invaders. Biological Invasions, 1(1), 3–19. https://doi.org/10.1023/A:1010034312781

Pimentel D, Mcnair S, Janecka J, Wightman J, Simmonds C, O’connell C, Wong E, Russel L, Zern J, Aquino T, Tsomondo T (2001) Economic and environmental threats of alien plant, animal, and microbe invasions. Ecosystems and Environment, 84(1), 1–20. https://doi.org/10.1016/S0167-8809(00)00178-X

Prado SDS, Lopes JRS, Demétrio CGB, Borgatto AF, Almeida RPD (2008) Host colonization differences between citrus and coffee isolates of *Xylella fastidiosa* in reciprocal inoculation. Scientia Agricola, 65(3), 251–258. https://doi.org/10.1590/S0103-90162008000300005

Purcell AH, & Finlay, A (1979) Evidence for Noncirculative Transmission of Pierce’s Disease Bacterium by Sharpshooter Leafhoppers. Phytopathology, 69(4), 393. https://www.apsnet.org/publications/phytopathology/backissues/Documents/1979Articles/Phyto69n04_393.PDF

Rathé AA, Pilkington LJ, Hoddle MS, Spohr LJ, Daugherty MP, Gurr GM (2014) Feeding and development of the glassy-winged sharpshooter, *Homalodisca vitripennis*, on Australian native plant species and implications for Australian biosecurity. PLOS ONE, 9(3), e90410. https://doi.org/10.1371/JOURNAL.PONE.0090410

Redak RA, Purcell AH, Lopes JRS, Blua MJ, Mizell III RF, Andersen PC (2004) The biology of xylem fluid–feeding insect vectors of *Xylella fastidiosa* and their relation to disease epidemiology. Annual Review of Entomology, 49(1), 243–270. https://doi.org/10.1146/annurev.ento.49.061802.123403

Roques A, Fan JT, Courtial B, Zhang YZ, Yart A, Auger-Rozenberg MA, Denux O, Kenis M, Baker R, Sun JH (2015) Planting sentinel European trees in Eastern Asia as a novel method to identify potential insect pest invaders. PLoS ONE, 10(5), 1–19. https://doi.org/10.1371/journal.pone.0120864

Santini A, Ghelardini L, De Pace C, Desprez-Loustau ML, Capretti P, Chandelier A, Cech T, Chira, D, Diamandis S, Gaitniekis T, Hantula J, Holdenrieder O, Jankovsky L, Jung T, Jurc D, Kirisits T, Kunca A, Lygis V, Malecka M, … Stenlid J (2013) Biogeographical patterns and determinants of invasion by forest pathogens in Europe. New Phytologist, 197(1), 238–250. https://doi.org/10.1111/j.1469-8137.2012.04364.x

Saponari M, Boscia D, Altamura G, Loconsole G, Zicca S, D’Attoma G, Morelli M, Palmisano F, Saponari A, Tavano D, Savino VN, Dongiovanni C, Martelli GP (2017) Isolation and pathogenicity of *Xylella fastidiosa* associated to the olive quick decline syndrome in southern Italy. Scientific Reports, 7(1), 1–13. https://doi.org/10.1038/s41598-017-17957-z

Saponari M, D’Attoma G, Abou Kubaa R, Loconsole G, Altamura G, Zicca S, Rizzo D, Boscia D (2019) A new variant of *Xylella fastidiosa* subspecies multiplex detected in different host plants in the recently emerged outbreak in the region of Tuscany, Italy. European Journal of Plant Pathology, 154(4), 1195–1200. https://doi.org/10.1007/s10658-019-01736-9

Saponari M, Giampetruzzi A, Loconsole G, Boscia D, Saldarelli P (2019) *Xylella fastidiosa* in olive in apulia: Where we stand. Phytopathology, 109(2), 175–186. https://doi.org/10.1094/PHYTO-08-18-0319-FI

Scally M, Schuenzel EL, Stouthamer R, Nunney L (2005) Multilocus sequence type system for the plant pathogen *Xylella fastidiosa* and relative contributions of recombination and point mutation to clonal diversity. Applied and Environmental Microbiology, 71(12), 8491–8499. https://doi.org/10.1128/AEM.71.12.8491-8499.2005

Schaad NW, Postnikova E, Lacy G, Fatmi M, Chang CJ (2004) Xylella fastidiosa subspecies: *X. fastidiosa* subsp *piercei*, subsp. nov., *X. fastidiosa* subsp. *multiplex* subsp. nov., and *X. fastidiosa* subsp. *pauca* subsp. nov. Systematic and Applied Microbiology, 27(3), 290–300. https://doi.org/10.1078/0723-2020-00263

Simberloff D, Martin JL, Genovesi P, Maris V, Wardle DA, Aronson J, Courchamp F, Galil B, García-Berthou E, Pascal M, Pyšek P, Sousa R, Tabacchi E, Vilà M (2013) Impacts of biological invasions: what’s what and the way forward. Trends in Ecology & Evolution, 28(1), 58–66. https://doi.org/10.1016/J.TREE.2012.07.013

Soliman T, Mourits MCM, van der Werf W, Hengeveld GM, Robinet C, Lansink AGJMO (2012) Framework for modelling economic impacts of invasive species, applied to pine wood nematode in Europe. PLOS ONE, 7(9), e45505. https://doi.org/10.1371/JOURNAL.PONE.0045505

Steiner G and Buhrer EM (1934) Aphelenchoides xylophilus n. sp., a nematode associated with blue-stain and other fungi in timber. Journal of Agricultural Research, 48(10), 949–951. https://naldc.nal.usda.gov/download/IND43968533/PDF

Stergiopoulos L and Gordon TR (2014) Cryptic fungal infections: The hidden agenda of plant pathogens. Frontiers in Plant Science, 5, 506. https://doi.org/10.3389/FPLS.2014.00506/BIBTEX

Strona G, Carstens CJ, Beck PSA (2017) Network analysis reveals why *Xylella fastidiosa* will persist in Europe. Scientific Reports, 7(1), 1–8. https://doi.org/10.1038/s41598-017-00077-z

Tomoshevich M, Kirichenko N, Holmes K, Kenis M (2013) Foliar fungal pathogens of European woody plants in Siberia: an early warning of potential threats? Forest Pathology, 43(5), 345–359. https://doi.org/10.1111/efp.12036

Vettraino AM, Roques A, Yart A, Fan JT, Sun JH, Vannini A (2015) Sentinel trees as a tool to forecast invasions of alien plant pathogens. PLoS ONE, 10(3), 1–15. https://doi.org/10.1371/journal.pone.0120571

Vettraino AM, Santini A, Nikolov C, Grégoire JC, Tomov R, Orlinski A, Maaten T, Sverrisson H, Økland B, Eschen R (2020) A worldwide perspective of the legislation and regulations governing sentinel plants. Biological Invasions, 22(2), 353–362. https://doi.org/10.1007/S10530-019-02098-3/FIGURES/2

Walther GR, Roques A, Hulme PE, Sykes MT, Pyšek P, Ku□hn I, Zobel M, Bacher S, Botta-Dukát Z, Bugmann H, Czúcz B, Dauber J, Hickler T, Jarošík V, Kenis M, Klotz S, Minchin D, Moora M, Nentwig W, … Settele J (2009) Alien species in a warmer world: risks and opportunities. Trends in Ecology and Evolution, 24(12), 686–693. https://doi.org/10.1016/j.tree.2009.06.008

Wells JM, Raju BC, Hung H-Y, Weisburg WG, Mandelco-Paul L, Brenner DJ (1987) *Xylella fastidiosa* gen. nov., sp. nov: Gram-negative, xylem-limited, fastidious plant bacteria related to *Xanthomonas* spp. International Journal of Systematic Bacteriology, 37(2), 136–143. https://doi.org/10.1099/00207713-37-2-136

XF-ACTORS (2017) Screening of olive cultivars for searching sources of resistance to *Xylella fastidiosa*. https://www.xfactorsproject.eu/screening-cultivars-resistance-xf/ [Accessed on 06.06.2022]

Yuan X, Morano L, Bromley R, Spring-pearson S, Stouthamer R, Nunney L (2010) Multilocus sequence typing of *Xylella fastidiosa* causing Pierce’s disease and oleander leaf scorch in the United States. Phytopathology, 100(6), 601–611. https://doi.org/10.1094/PHYTO-100-6-0601

Zhao BG, Futai K, Sutherland JR, Takeuchi Y (2008) Pine wilt disease. Springer (ed.), Tokyo Japan, Vol. 17, p. 459. https://link.springer.com/book/10.1007/978-4-431-75655-2?noAccess=true

