## supplementary file 1 for "Measuring the threat from a distance: insight into the complexity and perspectives for implementing sentinel plantation to test host range of *Xylella fastidiosa*"

### Administrative burdens and delays, a comparative view

#### 1. A first attempt in Apulia (Italy) – full procedural pathway

a) May 2016 – Colleague contacted at CNR Lecce in order to establish collaborative links and locate a suitable location for the nursery. Location is found and experimental design is finalized.

b) 24/10/2016 - E-mail from JC Grégoire to *personincharge1*@politicheagricole.it, applying for a permit – no reply

c) Exchanges with CNR Lecce colleague regarding the drafting of an official letter to the Italian and Apulian authorities

d) 3/11/2016 – JC Grégoire sends letter to the Italian and Apulian authorities, following colleague's model

e) 3/2/2017 – Claude Bragard and JC Grégoire send modified version of the letter to the Apulian authorities (Dott. *personincharge2*)

f) 4/2/2017 – CNR Lecce colleague indicates that the request should be written by an Italian partner.

g) 6/2/2017 – C Bragard contacts Italian partner, a nursery owner involved in the EU proposal "LUB-XYL".

h) 8/2/2017 – C Bragard sends request to the Apulian authorities (Dott. *personincharge3*)

i) 4/4/2017 – reply (very short and rather obscure) from *personincharge4* (Regione Puglia):

   *Avviare le procedure di cui all'art. 8 del DM 7/12/2016 (ex art. 5, Decisione 2015/789 s.m.i.)ai fini dell'autorizzazione, in deroga al divieto di impianto delle piante ospiti ed in conformità alle condizioni di cui al Titolo X del d.lgs. 214/2005, per fini scientifici.*

j) 3/5/2017 – Explanations from Alberto Santini: list of criteria to be fulfilled:
- a) name and address of the responsible of the activity;
- b) scientific name of the plant material;
- c) type of material (seedlings, adult plants, cuttings, etc.)
- d) quantity of the material
- e) origin and provenance of the material;
- f) length, kind and aims of the research and, at least, a short summary of the activities;
- g) address and description of the quarantine site;
- h) plantation site (once you have the permit);
- i) destruction method, if requested.
- j) point of entrance if plant material is imported from third countries. (not in our case)

k) 5/5/2017 – New request from the Belgian group

l) 11/5/2017 – Authorization from the Apulian authorities.

m) 15/2/2018 – Plantations planned around mid-February

**Total duration of the process: 21 months…**

#### 2. Second attempt: Majorca (Spain)

a) 26.02.2018 - Request to the Vice-chancellor at Universitat de les Illes Balears (UIB)

b) 06.3.2018 - Reminder of the request

c) 06.3.2018 - Reply from Vice-chancellor saying we need to ask permit to the local Government (Conselleria de Medi Ambient, Agricultura i Pesca)

d) 16.3.2018 - Permit and approval sent from local government (Conselleria)

e) 20.3.2018 - Meeting of Rector and Vice-chancellors at UIB

f) 21.03.2018 - Communication of the approval for the sentinel plantation at UIB

g) 30.03.2018 – Plantation begins

Total duration of the process: slightly more than 1 month…

#### 3. To summarise

- Complexity and diversity of the administrative procedure.
- Absolute need to benefit from strong, organised and well-informed local partnership.
